# Fungal Mst3-proteins are involved in fungal innate immunity needed for the recognition of bacteria surrounding the hyphae, as well as for plant pathogenicity

**DOI:** 10.1101/2025.11.25.690610

**Authors:** Shanshan Gong, Xiaorong Lin, Shenghua Liu, Stefan Olsson, Guodong Lu, Zonghua Wang, Ya Li

**Affiliations:** State Key Laboratory of Agricultural and Forestry Biosecurity, College of Plant Protection, Fujian Agriculture and Forestry University, Fuzhou 350002, China; Synthetic Biology Center, College of Future Technologies, Fujian Agriculture and Forestry University, Fuzhou, China; Institute of Oceanography, Minjiang University, Fuzhou, China

**Keywords:** Innate immunity, MAMP, PAMP, inflammasome, bacteria, NLR, ROS

## Abstract

Fungal innate immunity overall responses are as fast as and resemble mammalian innate immunity. It does not, however, employ membrane toll-like receptors (TLRs), but should employ endocytosis of non-fungal molecular patterns recognized by nuclear-localizing receptors (NLR). Downstream, both types of receptors in mammals are Mammalian Ste20 kinases (MSTs). We identified single MST3 orthologs in the plant pathogens *Fusarium graminearum* (*FgMST3*) and *Magnaporthe oryzae* (*MoMST3*). We knocked out both genes and investigated mutants using a standard panel of tests for growth, development, and pathogenicity for each respective fungus. Both *ΔFgMST3* and *ΔMoMST3* strains showed reduced pathogenicity. The deletions negatively affected conidia production and conidia germination but had little effect on growth rate. However, the two mutants reacted differently to some stress treatments, such as Zn^2+^ and gentamicin. Furthermore, we constructed an innate immunity reporter system for *F. graminearum* based on a gene with fast responses to bacterial MAMPs to detect less than 4-hour responses to non-self-molecular patterns (NSMP) like bacterial outer membrane vesicles (OMVs) as well as trace levels of sucrose, as a plant NSMP. The reporter gene responses to OMVs of MST3 mutant strains were severely reduced. Our results suggest that both MoMst3 and FgMst3 are involved in fungal innate immunity downstream of unknown NLR proteins, motivating studies to try to identify genes for the NLR-like receptors. Finding such and investigating how they work and vary between fungal species and strains should be useful for understanding fungal biotic interactions with viruses, bacteria, plants, and animals.

## Introduction

Sterile20 kinases (Ste20) are named after a yeast kinase that, when deleted, made yeast sterile (1). A subclass of Ste20 kinases is Mammalian-like STe20 kinases (MSTs). These proteins are involved in non-self-signaling relaying signals from Toll-Like-Receptors (TLRs) on the cell membrane, or intracellularly from Nuclear Localizing Receptors (NLRs) (2, 3). In mammals, the MSTs fall into two groups of GCKII and GCKIII, containing MST1/2 and MST3, respectively, depending on their structure (3). Outside the MST functioning Ste20 kinases, other Ste20 kinases can have roles in cell volume sensing and regulation of Cl-transport. Yeast Ste20, for example, controls a cell shrinkage-activated MAPK cascade that regulates organic osmolyte accumulation (2).

Fungi have no identified TLRs (4) but show an overall mammalian-like innate immunity response (5) that differs from plant innate immunity, possibly reflecting that in the early eukaryote evolution, plants split off from fungi-metazoans (6) and innate immunity is present already in Protista (7, 8). Since innate immunity also employs Nuclear Localized Receptors (NLRs), these could be the receptors fungi use to recognise bacteria. NLRs are possibly evolutionary older than TLRs and occur already in the first unicellular Eukaryotes (7, 8), and seem to be especially important in epithelia naturally colonized by bacteria, such as in the gut (9–11). In plants, innate immunity seems different in different tissues, and the most studied TLR-triggered immunity is very limited in roots or the root epithelium, while NLRs are present (12, 13). The innate immunity proteins in fungi recognizing microbial-associated molecular patterns are thus most likely NLRs of the STAND type that are similar to proteins in the inflammasome and probably involve other repeats than just LRRs, like, for example, WD40 repeats (approx. 40 aa repeats ending with the amino acid WD, InterPro) (14–17).

Considering all the above, orthologous fungal MST kinases, especially of the MST3 type, could be involved in non-self-signaling downstream of fungal NLR-like non-self-recognizing molecular pattern (NSMPs) recognition receptors, triggering innate immunity responses and influencing fungal pathogenicity and interactions with bacteria. As with bacteria, the host plant is also a non-self-entity that sheds molecular patterns into the environment just outside the plant cells, into the phyllosphere/apoplast/rhizosphere, that are different from what is present in fungi. Such plant NSMPs could be plant hormones, cellulose oligomers, pectin oligomers, sucrose, and other fungi NSMPs that can be taken up through endocytosis and be presented to NLR-type receptors and trigger inflammasome-like proteins (18). In line with this, sucrose not generally present in soil except in the inner rhizosphere and the plant apoplast outside the plant cells, has been shown to stimulate innate immunity-like responses in a *F. graminearum* (5), and it is present in the first plant contact zones for infecting fungi. Thus, endocytosed plant NSMPs of different origins might be recognized by hitherto not identified fungal NLR-like NSMP-receptors that are, as in animals, dependent on MST3 for downstream signaling (2, 3).

We selected 2 relatively closely related fungi, *Fusarium graminearum* and *Magnaporthe oryzae,* with different lifestyles to investigate if MST3 orthologs are involved in pathogenesis. Both fungi are well-studied model pathogen fungi with established and efficient gene transformation protocols (19). *F. graminearum* has to be able to handle bacteria on plant surfaces both above ground and especially below ground in the seedling rhizosphere (20), while *M. oryzae* mainly meets bacteria in the phyllosphere during conidia germination (21). Both fungi first infect as biotrophs and then, probably because of stresses to the fungus inside the plant, switch to a necrotrophic lifestyle (19). In the biotrophic/necrotrophic transition, plant innate immunity is strongly induced. Despite this, the pathogen normally copes and even enhances plant cell death, killing the plant cells and creating necroses, in compatible interactions. In the preceding biotrophic stage, plant innate immunity is suppressed by the fungus (22). Consequently, fungal innate immunity signaling should be most important to the fungus in the biotrophic establishment stage both to counteract interfering bacteria and to identify that it has entered a plant to turn on fungal genes needed to establish the interaction with the plant, and suppress plant innate immunity until the fungus has biomass and resources enough to cope with a switch to necrotrophy with accompanying increased plant defences (22).

We identified an MST3-like putative protein in each of the two fungi and could show that such MST3 orthologs can be found from Archaea across all investigated Eukaryotes. The deletion of both these MST3-kinases had affected growth, conidia production, conidia germination, stress phenotypes, and plant pathogenicity in the respective fungus. Furthermore, we found that the deletions had negative effects on fungal innate immunity responses triggered by bacterial NSMPs in F. graminearum were we had an established system for exposing the fungus to bacterial NSMPs. This indicates that MST3 signaling is important for fungal recognition of bacteria that might be pathogenic. It might also be important for sensing plant-produced NSMPs, signaling to the fungus that it is in the inner rhizosphere of a plant root or on a plant surface.

## Results

### Bioinformatics search for a putative *MST3* gene in the genomes of both fungi

In Rice, the BSR1 gene, encoding a kinase, works downstream of TLR receptors to relay signals in innate immunity (23) and might do the same downstream of NLR receptors. Thus, we looked for a similar protein in *F. graminearum* and found one. We also found that the closest human homologue to rice OsBsr1 is HsIrak4, which has a similar role in mammals, but both OsBsr1 and HsIrak4 are too short to have the same function as the closest homologue in *F. graminearum*. We then did a protein blast search in animal protein sequences at NCBI to search for a homologue to the protein sequence we had found in *F. graminearum*. We looked especially for experimentally well-described mammalian proteins since fungal innate immunity responses have similarities with innate immunity in mammals (5). It was then that we realized that the *F. graminearum* gene could encode a well-conserved eukaryotic Mst3 kinase involved in relaying signals from NLRs to trigger innate immunity. The *F. graminearum* BSR1-like protein used as bait for a BLAST against human sequences gave one good candidate that is known as an Mst3 kinase in animals (E-value 5 x E-127. 67% identities and 82% positives and no gaps) so we named our bait gene FgMst3. In *M. oryzae* there is an almost identical predicted protein to the FgMst3 (E-value 0.0, 62% identities and 71% positives and 12% gaps) and we named that MoMst3. Since Mst3 kinases are conserved in both fungi and mammals we downloaded potential orthologuos protein sequences for a range of organisms at further and further evolutionary distances from our two fungi and used the rice BSR1 and human Irak4 that are known to relay signals from TLR receptors as an outgroup.

All the putative Mst3-like proteins from Archaea to *Homo sapiens* we found can potentially be involved in relaying signals from NLR receptors in innate immunity, since innate immunity is evolutionarily ancient and appears to have evolved at the dawn of eukaryotes to recognize bacteria invading their cytoplasm (7) (**Fig. 1**).

**Figure 1.**
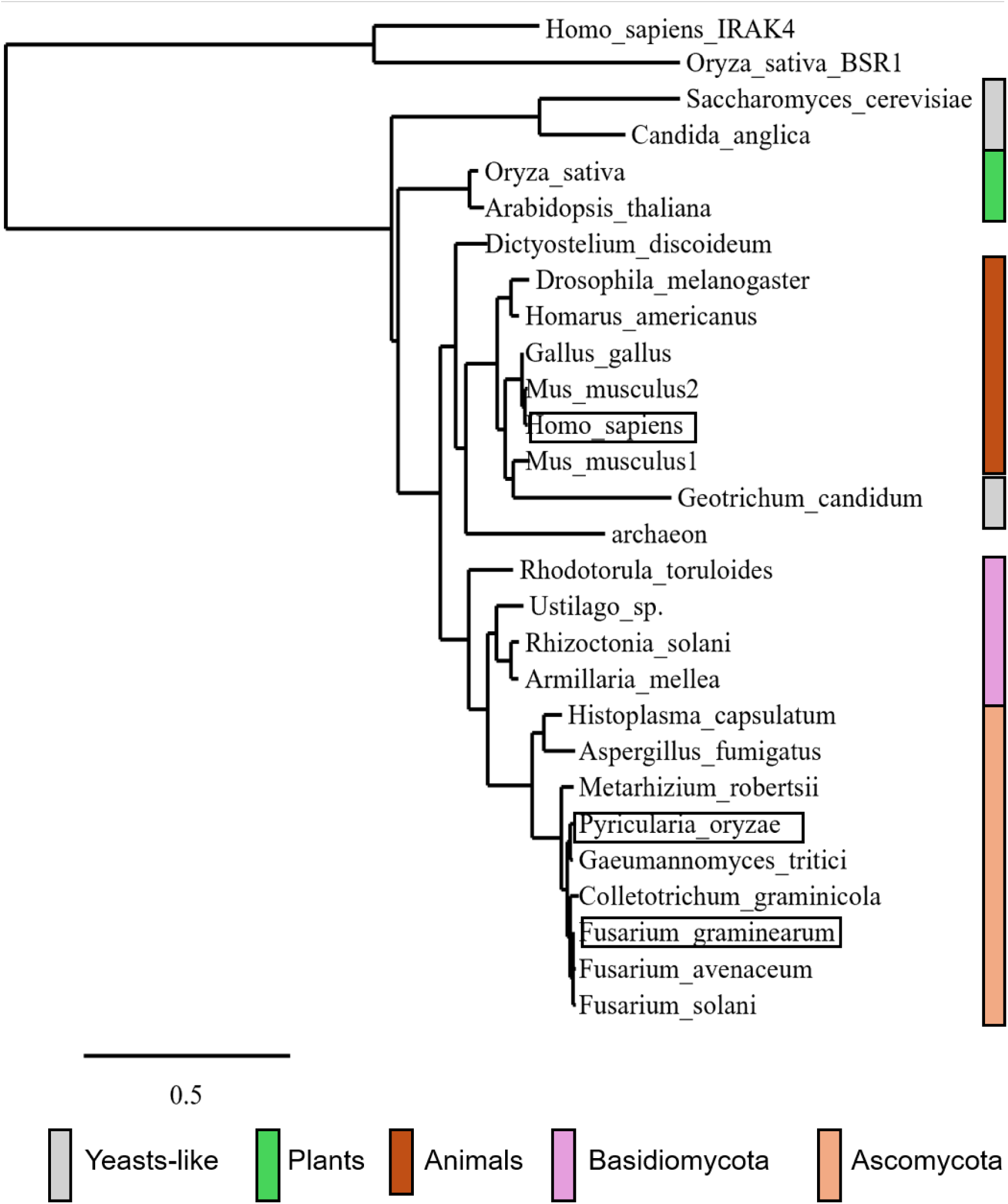
Phylogeny of MST3 protein homologues. The mammalian proteins are the 2 canonical MST3s from mouse and the human MST3 (49). The names *F. graminearum* and *Pyricularia oryzae* (the teleomorph equivalent of *Magnaporthe oryzae*), as well as the human MST3 protein they are compared with, are marked with black boxes.

The Mst3-like kinases are different from BSR1 and IRAK4 kinases that are known to relay signals from TLR cell surface receptors (23, 24) and appear truncated to only the N-terminal kinase part of the Mst3-like proteins (**Fig. S1** Alignment of the proteins in the phylogeny)

The mammalian Mst3 proteins are known to have the kinase domain in the first half of the protein and both nuclear localization signal (NLS) and a possible nuclear export signal (NES) in the C-terminal end. In between the NLS and the NES is a caspase cleavage site for caspase proteases in mammals that can be activated during apoptosis (25). For fungi, it should be a metacaspase cleavage site since fungal metacaspases cleave similarly to the main “executioner” caspase, caspase 3 (26, 27).

We used a prediction website to predict caspase/metacaspase cleavage sites (https://scap.cbrc.pj.aist.go.jp/ScreenCap3/index.php for caspase cleavage) and alignment of the known NLS signal in the human protein to predict the relative p[sition position of putative NLS signals in the fungal sequences (**Fig. 2**, and see **Supplemental File SF1** for the more detailed bioinformatics done)

**Figure 2.**
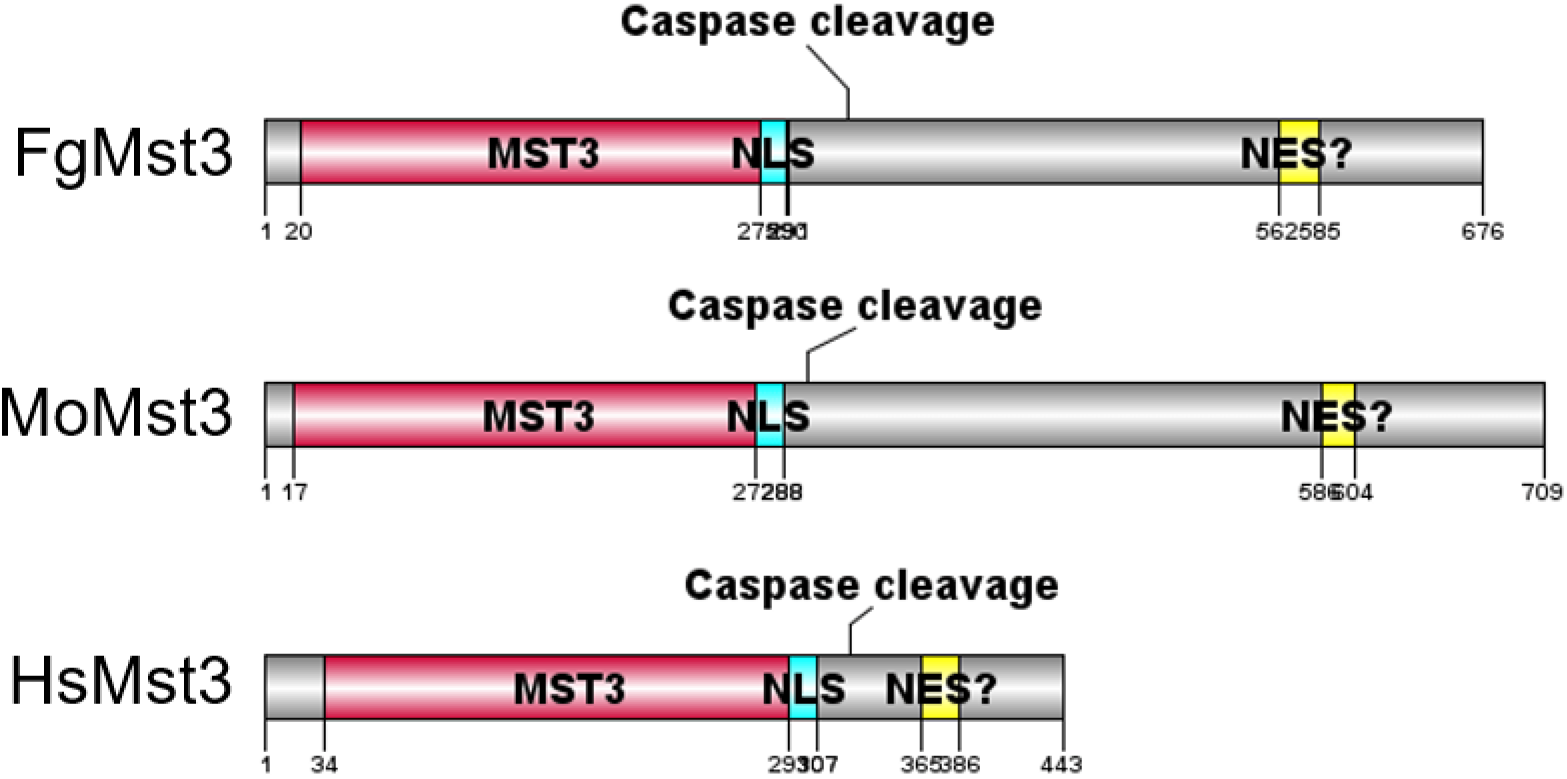
Predicted domains and cleavage sites in the 2 fungal proteins and compared with known domains and the cleavage sites in human Mst3 (HsMst3). NLS=nuclear localization signal. NES?= suggested nuclear export signal region based on a conserved alignment of the two fungal proteins with the 335-385 section of the HsMst3 implicated to contain a NES.

The shorter C-terminal of the human protein contains the NES signal that is less well defined compared to the NLS, so we aligned the truncated C-terminal after the predicted caspase cleavage site. In conclusion, cleavage by caspase at the first cleavage site of all 3 proteins should cleave off an NES signal, leading to a caspase-mediated nuclear localization of the fungal MST3 during apoptosis. There is, however, a large extra chunk of approximately 200 AA, just after the predicted caspase cleavage site that is conserved in the fungal proteins, implying that these fungal Mst3 proteins could have additional functions in the cytoplasm after being cleaved off, but maybe also before cleavage.

### Deletion of genes in both fungi and the growth effects of the deletions on the growth and development of conidia

The putative *FgMST3* and *MoMST3* genes were deleted successfully, and the resulting *ΔFgmst3* and *ΔFgMST3* strains were checked for possible ectopic integrations of the HPH gene used as a selection marker for deletions using GO-qPCR (see Materials and Methods). We constructed both *FgMST3-GFP* and *MoMST3-GFP* strains from these respective mutant strains (**Fig. S2, S3, S4**). As extra control, we used RT-qPCR to show that no extra working copies of the two MST3 genes have been created or are present in the genome (**Fig. S5).** We could also show that both FgMst3-GFP and MoMst3-GFP are mainly localized in the cytoplasm of the strains grown *in vitro,* as expected since the Mst3 proteins have intact putative NES signals (**Fig. S6 and Fig. S7**).

**Figure 3.**
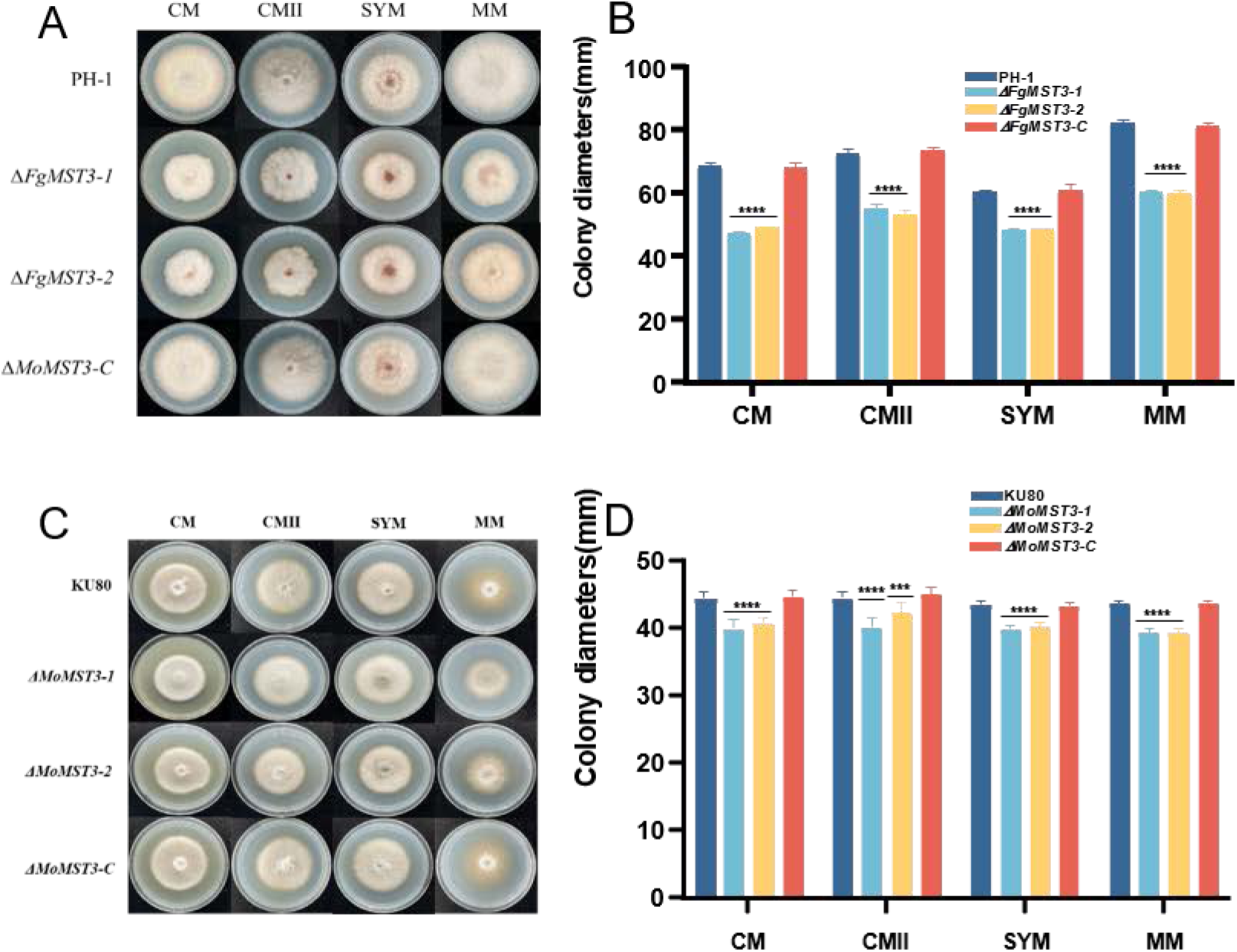
Growth effects of *MST3* deletions on CM, CMII, SYM, and MM media. (A and C) Colony morphology (B and D) and colony diameter after 3- or 7-day incubation of *F. graminearum* or *M. oryzae* cultures, respectively. Error bars in (B and D) show SEMs. 3 stars indicate a P_same_ <0.001 and 4 stars indicate a P_same_ <0.0001 probability for the null hypothesis that these measurements are the same as for the respective background strain.

**Figure 4.**
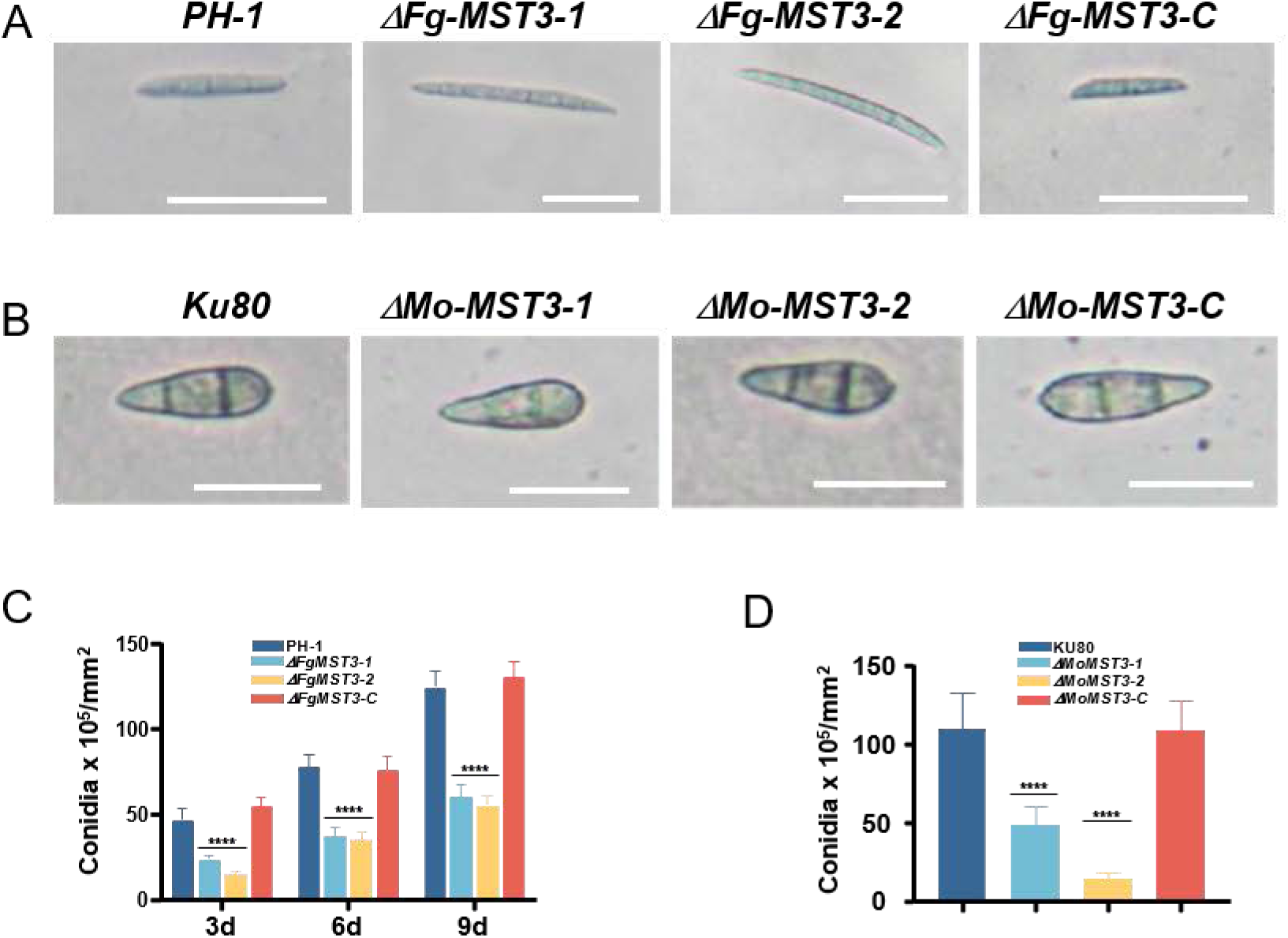
***F. graminearum* and *M. oryzae* conidia morphology and conidia production** Conidia morphology (A) and (B), and conidia production in CMC liquid medium (C) and (D). Error bars show SEMs, and 4 stars indicate P_same_ <0.0001 probability for the null hypothesis that these measurements are the same as for the respective background strains.

Both mutant strains have a slower colony radial growth rate than their respective background strains, *PH-1* and *KU80,* respectively, on all 4 media tested. The growth rate of *ΔFgMST3* strains decreased more than that of *ΔMoMST3* strains, compared to their respective background strains (**Fig. 3**).

Conidiation in both fungi was strongly affected by the deletions of *FgMST3* and *MoMST3* genes. Both mutant strains produced fewer conidia than their respective background strains and complementation strains. In addition, the conidia in *ΔFgMST3* strains appeared a bit longer than conidia from *PH-1,* while there was no difference between *ΔMoMST3* conidia and those from *KU80* (**Fig. 4**).

### The effect of *MST3* gene deletions on a panel of important fungal characteristics that can be tested *in vitro*

Conidial germination and rate of germination, as well as the tolerance of the fungal strains to stress treatments of the kind that a pathogenic fungus meets inside a plant, are such characteristics. In addition, there is a well-established appressorium formation assay available for *M. oryzae* (28).

Conidia of both the *ΔFgMST3* and the *ΔMoMST3* strains germinated well **(Fig. 5 A-D)** but faster than conidia from either PH-1 or KU80 and reached the same level of germination. This faster increase in germination per day was for *ΔFgMST3* strains between hours 2-4h and 6-8h for *ΔMoMST3* strains **(Fig. 5B and Fig. 5D)**. Appressorium formation from the conidia of *ΔMoMST3* strains was always lower from 4 h to 12 h (**Fig. 5E**).

**Figure 5.**
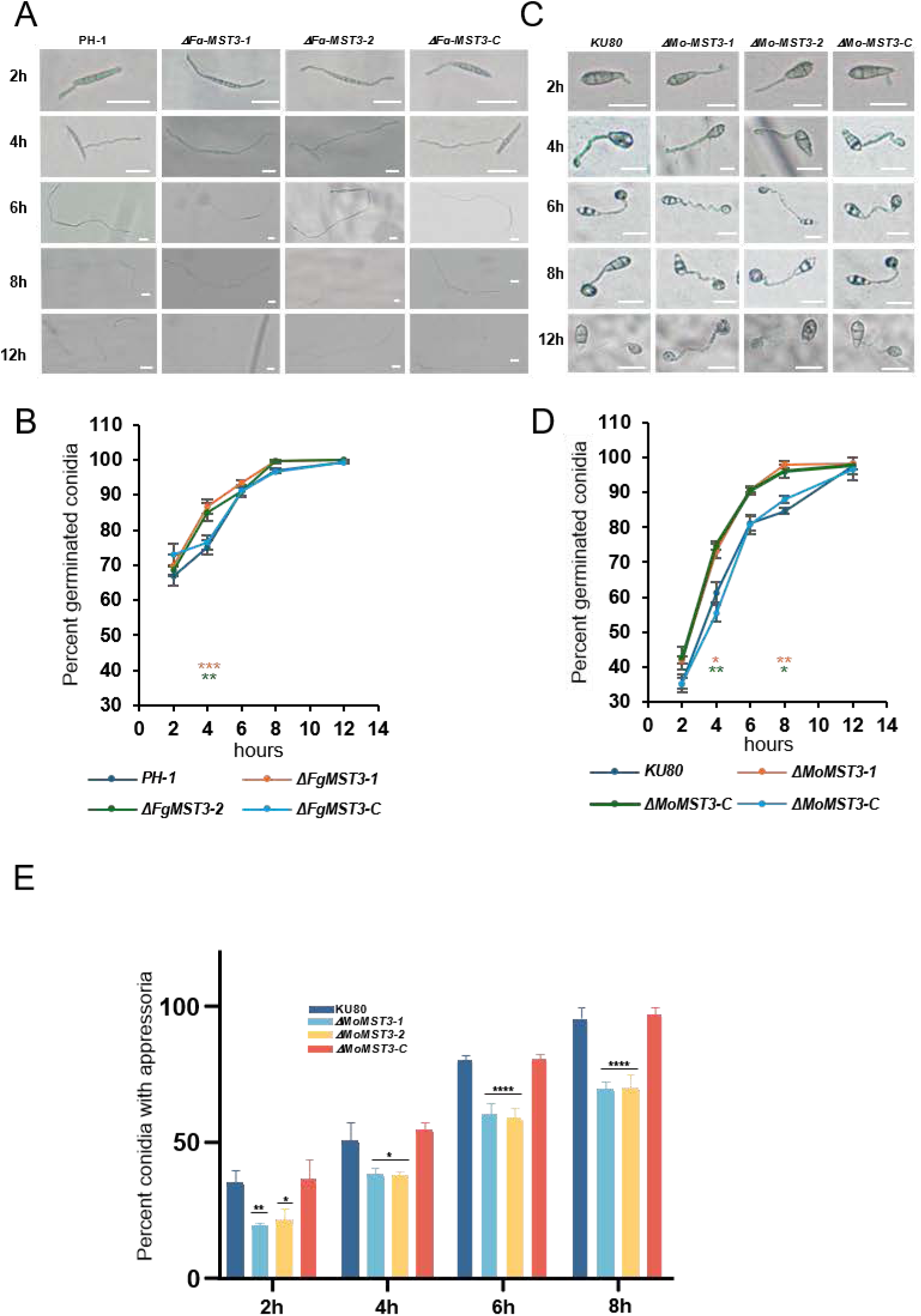
Effect of *FgMST3* and *MoMST3* mutations on conidia germination and conidia germination dynamics. Conidia germination morphologies after different times of germination (A and C), cumulative germination dynamics (B and D), and appressoria formation from the germinated conidia (E). Error bars indicate SEMs, and *, **, ***, and **** symbolize P_same_<0.05, <0.01, < 0.001, and <0.0001, respectively, for the null hypothesis that these measurements are the same for their respective background strains at the same timepoint.

A panel of standard stress treatment of relevance to the physiology of fungal plant pathogens were used to test whether the *ΔFgMST3* and the *ΔMoMST3* strains were affected in comparison to their background strains (**Fig. 6**). The H_2_O_2_ and SDS treatments both affect membrane stability, while the former also has effects on oxidizing DNA and proteins. CFW and CR treatments both have effects on cell wall integrity, while sorbitol and NaCl result in osmotic and osmotic/ionic stresses, respectively

**Figure 6.**
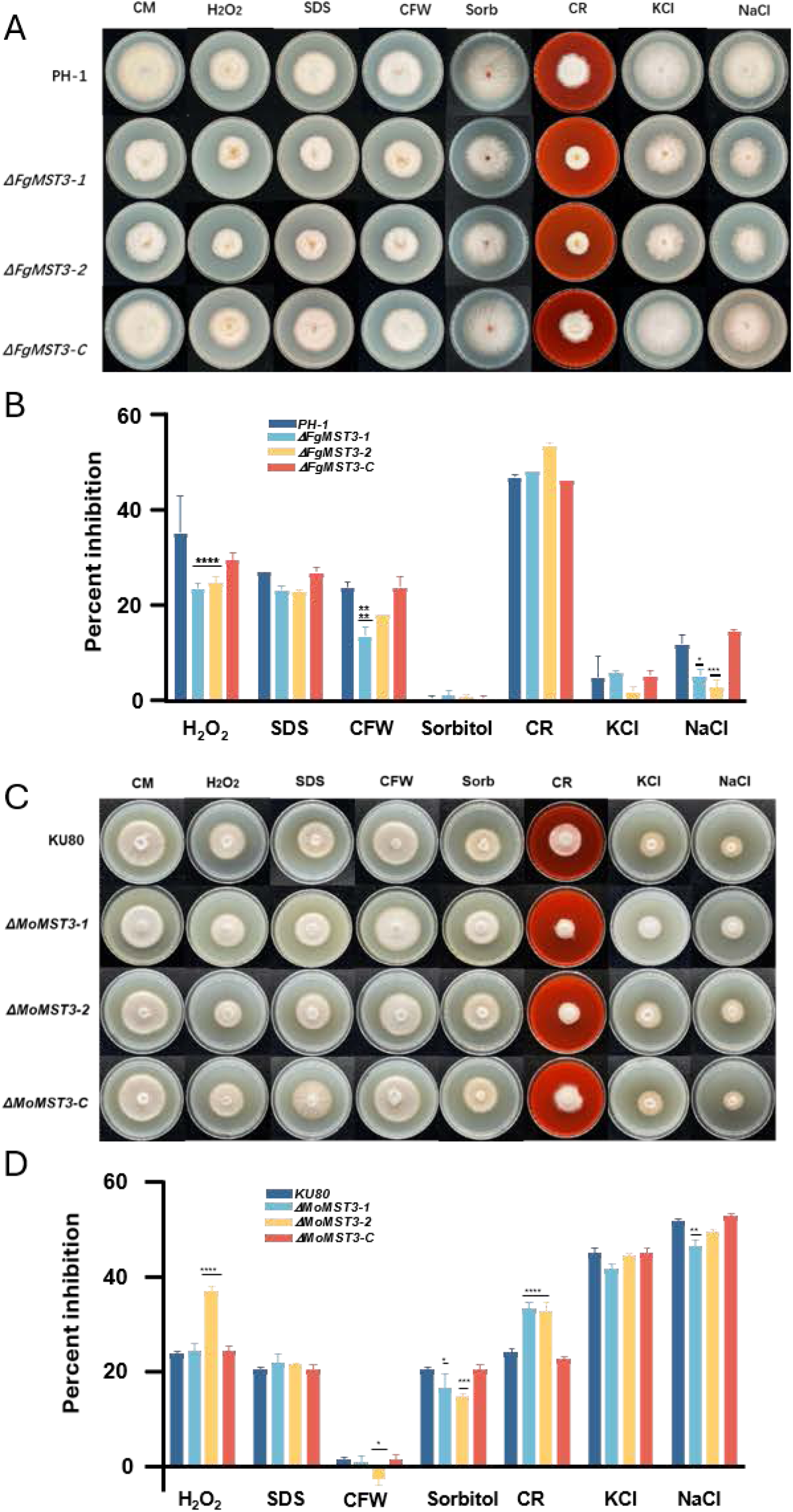
***F. graminearum and M. oryzae* strains grown on a panel of stress-inducing media** Colony morphology (A and C) on the different stress media and inhibition of colony diameter growth on the same media. The colony diameter for the background strains (PH-1 and Ku80) on CM without a stressing agent, minus the colony diameter on the stress medium, gives colony growth inhibition (B and D). The larger the value, the more inhibited by the stress medium. Error bars indicate SEMs, and *, **, ***, and **** symbolize P_same_<0.05, <0.01, < 0.001, and <0.0001, respectively, for the null hypothesis that these measurements are the same as for respective background strains.

Treatments with H_2_O_2_, SDS, and NaCl had less effect on the *ΔFgMST3* strain than on PH-1. Treatment with CR, on the other hand, was more inhibiting, at least for one of the *ΔFgMST3* strains, and NaCl had less effect on the *ΔFgMST3* strains **(Fig. 6B)**. Contrary to the *ΔFgMST3* strains, the *ΔMoMST3* strains were more affected by H_2_O_2_ than the control, thus less resistant, and they were also more strongly affected by CR (**Fig. 6D**).

Taken together, it appears that *M. oryzae* and *F. graminearum MST3* mutants react both similarly and differently to stresses, probably reflecting their respective biology as discussed in the discussion section.

### Effect of the *MST3*-deletion on the mutant strain’s pathogenicity

Both fungi have an initial biotrophic stage that is essential for the spread of the fungus inside the plant tissue (19). *F. graminearum* is mainly recognized as a problem when infecting wheat heads, causing head blight, but it also infects from soil, causing severe seedling blight (20). For *F. graminearum,* two standard pathogenicity assays were thus used, one for wheat coleoptiles and one for wheat heads. In both assays, the two *ΔFgMST3* strains showed strongly reduced pathogenicity. The coleoptile lesions became smaller for the *ΔFgMST3* strains (**Fig. 7 A-B**). In the wheat head assay, it was obvious that the main difference was a faster spread from the initial infected flower in the PH-1 through the rachis connecting the flowers and to other flowers and developing seeds (**Fig. 7 C-D**). Taken together, the deletion of the *FgMST3* gene negatively affected the *F. graminearum* pathogenicity on wheat.

**Figure 7.**
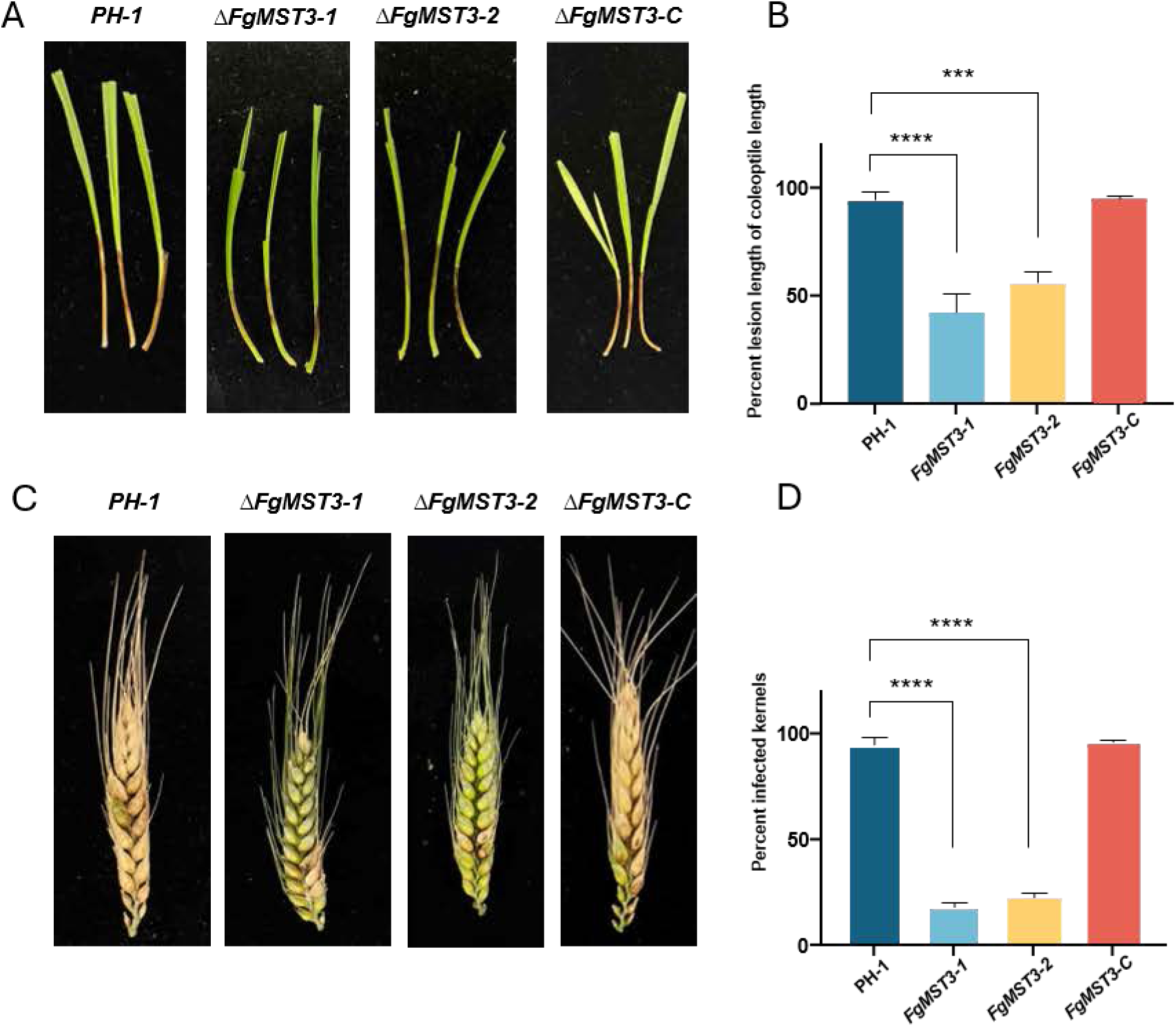
Pathogenicity test of *F. graminearum* strains on wheat coleoptiles and wheat heads. (A) Wheat coleoptiles with lesions at 7DPI (B) Average percent lesion length of coleoptile lengths below the point of inoculum at 7DPI. (C) Wheat heads at 14DPI after inoculation at the 3-4 flowers from the base of the head. (D) Average percent diseased spikelets at 14DPI. Error bars indicate SEMs, and *, **, ***, and **** symbolize P_same_<0.05, <0.01, <0.001, and <0.0001, respectively, for the null hypothesis that these measurements are the same as for the PH-1.

*M. oryzae* is a leaf pathogen of many grasses and can also infect wheat leaves from conidia germinating and forming an appressorium that penetrates the leaf epidermis, entering the plant and spreading between leaf cells as a biotroph. When the easily available soluble nutrients in a plant cytoplasm run out, the pathogen switches to the necrotrophic stage, killing the cell and degrading its components, using the released nutrients to form conidia, mediating the spread to new host plants. The *M. oryzae* necrotrophic activity creates lesions of plant leaf cells dying by a combination of fungal activities and the plant’s own defense, triggering programmed cell death (PCD) of the plant cells (29). The *ΔMoMST3* where less pathogenic than KU80 in all three assays performed **(Fig. 8 A-F**)

**Figure 8.**
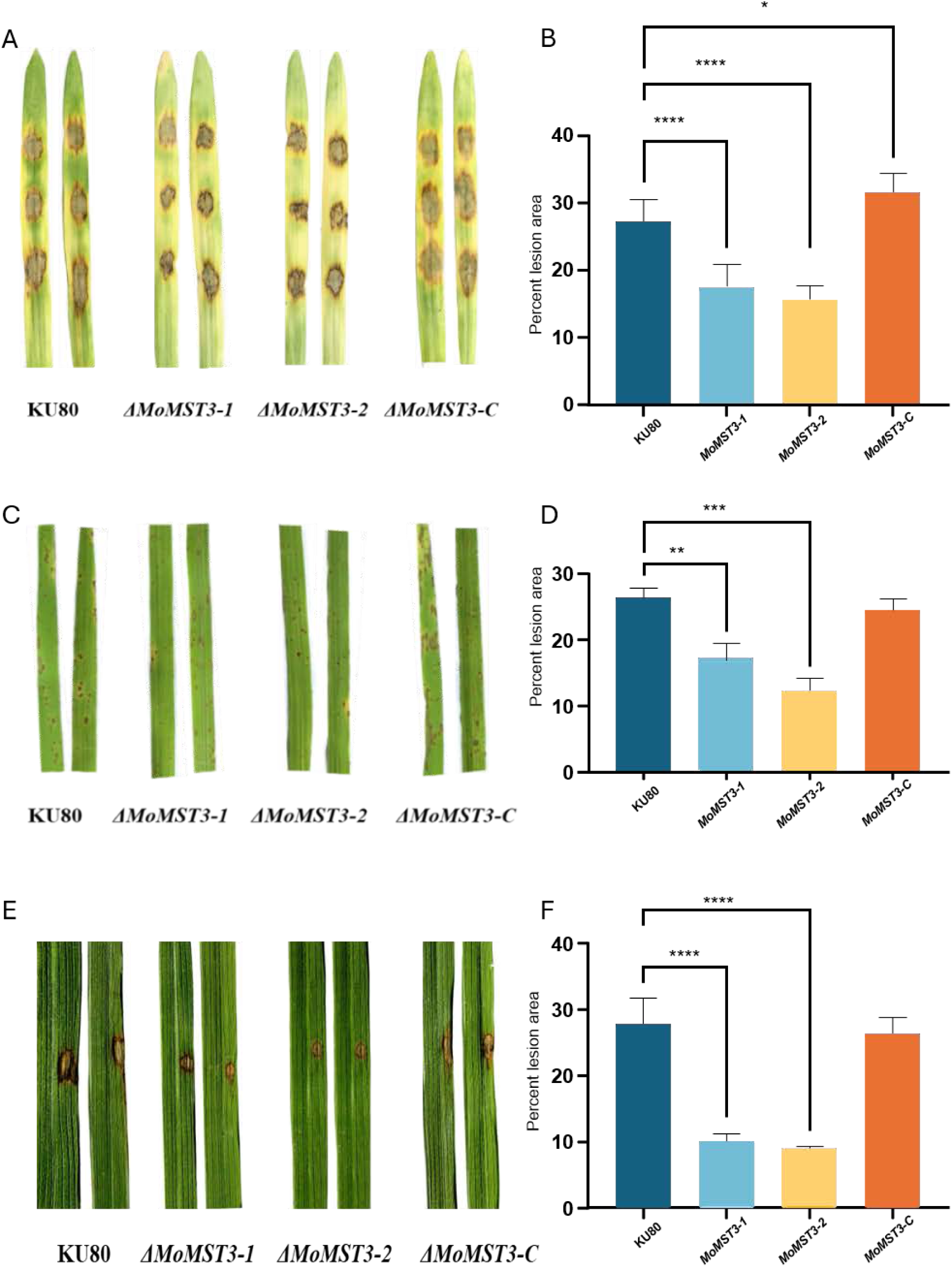
Lesion formation by *M. oryzae* strains on barley and rice leaves. Conidia were dripped on excised barley leaves (A-B). Lesion appearance on the leaves at 3-5DPI. (A) Average percent lesion area of available leaf area (B). Conidia were sprayed on excised rice leaves (C-D). Lesion appearance on the leaves at 5-7DPI. (C) Average percent lesion area of available leaf area (D). Conidia were added to press-injured spots on excised rice leaves (E-F). Lesion appearance on the leaves at 15DPI (E). Average percent lesion area of available leaf area (F). Error bars indicate SEMs, and *, **, ***, and **** symbolize P_same_<0.05, <0.01, <0.001, and <0.0001, respectively, for the null hypothesis that these measurements are the same as for the KU80.

### Additional stress treatments by Zn and Gentamicin

Uptake of both Zn and the weakly antifungal but strongly antibacterial compound gentamicin is dependent on the function of the fungal cell wall and cell membrane, as well as other processes in the fungus. Since the *ΔFgMST3* strains and the *ΔMoMST3* strains differed in their reaction to some of the stress treatments, and *M. oryzae* has melanized cell walls that *F. graminearum* lacks, we added assays for these stress treatments to investigate if these could show clearer differences between the deletion strains and their respective backgrounds. It turned out that *ΔFgMST3* strains were less sensitive (**Fig. 9 A, B**) than *ΔMoMST3* strains to Zn^2+^ stress (**Fig. 9 C, D**). The same applied to gentamicin addition. It even appears that gentamicin is not stressful at all to *F. graminearum,* as both the *PH-1* strain and the complement strain grow slightly better than controls without gentamicin. The largest positive effect of the gentamicin additions was for the two *MST3* mutant strains. **(Fig. 10 A-B and Fig. 10 C-D)**.

**Figure 9.**
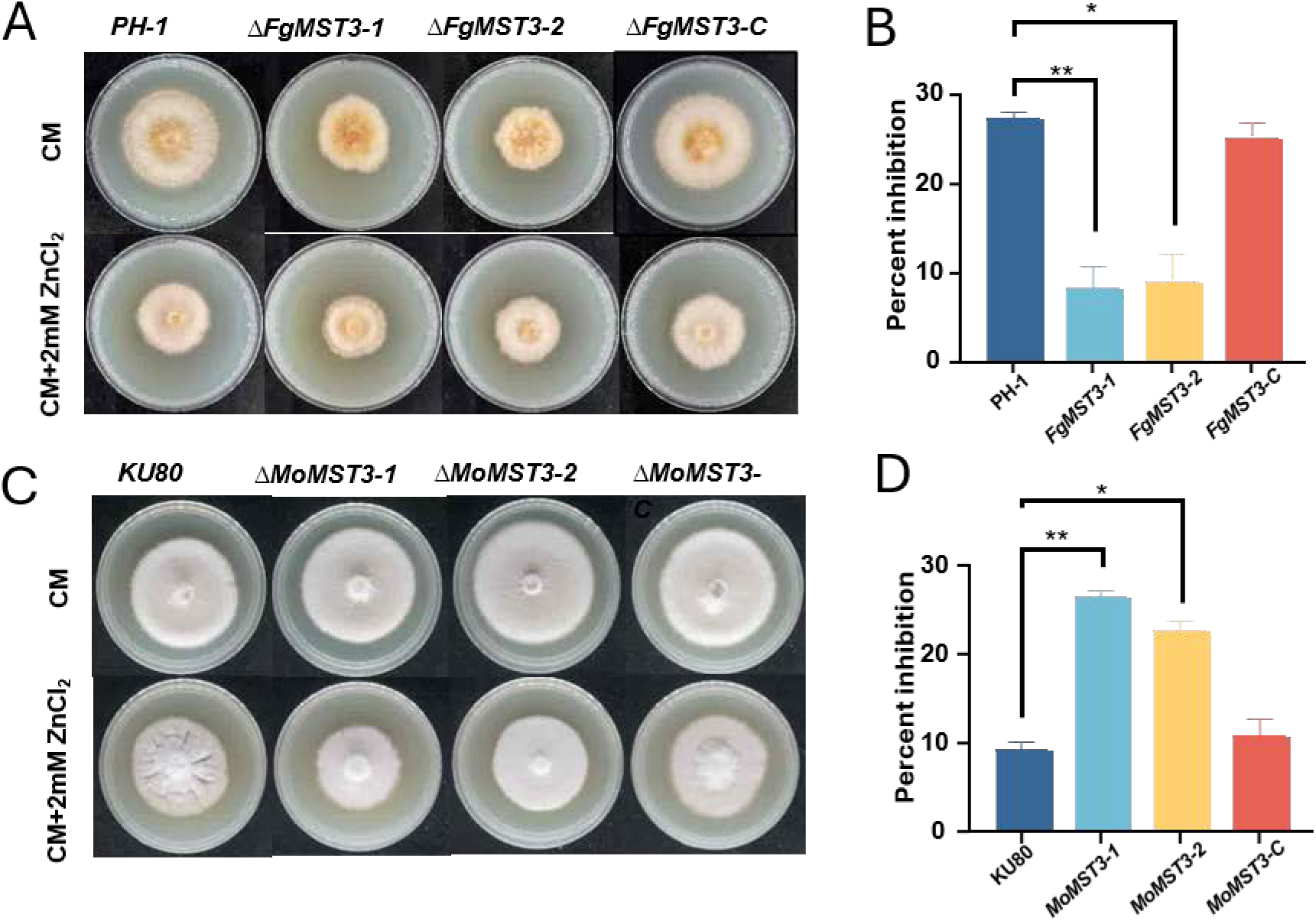
Inhibition of growth by Zn^2+^ for the different *F. graminearum* strains and *M. oryzae* strains. Colony morphology of *F. graminearum* strains in the stress medium (A) and inhibition of colony growth on the medium, colony diameter for PH-1 on control medium without stressing agent minus colony diameter on stress medium gives colony growth inhibition (the larger the value, the more inhibited by the stress medium) (B). Colony morphology of *M. oryzae* strains on the different stress media (C) and inhibition of colony growth on the same media, colony diameter for KU80 on control medium without stressing agent minus colony diameter on stress medium gives colony growth inhibition (the larger the value, the more inhibited by the stress medium (D). Error bars indicate SEMs, and *, **, ***, and **** symbolize P_same_<0.05, <0.01, < 0.001, and <0.0001, respectively, for the null hypothesis that these measurements are the same as for the PH-1 or KU80 strains.

**Figure 10.**
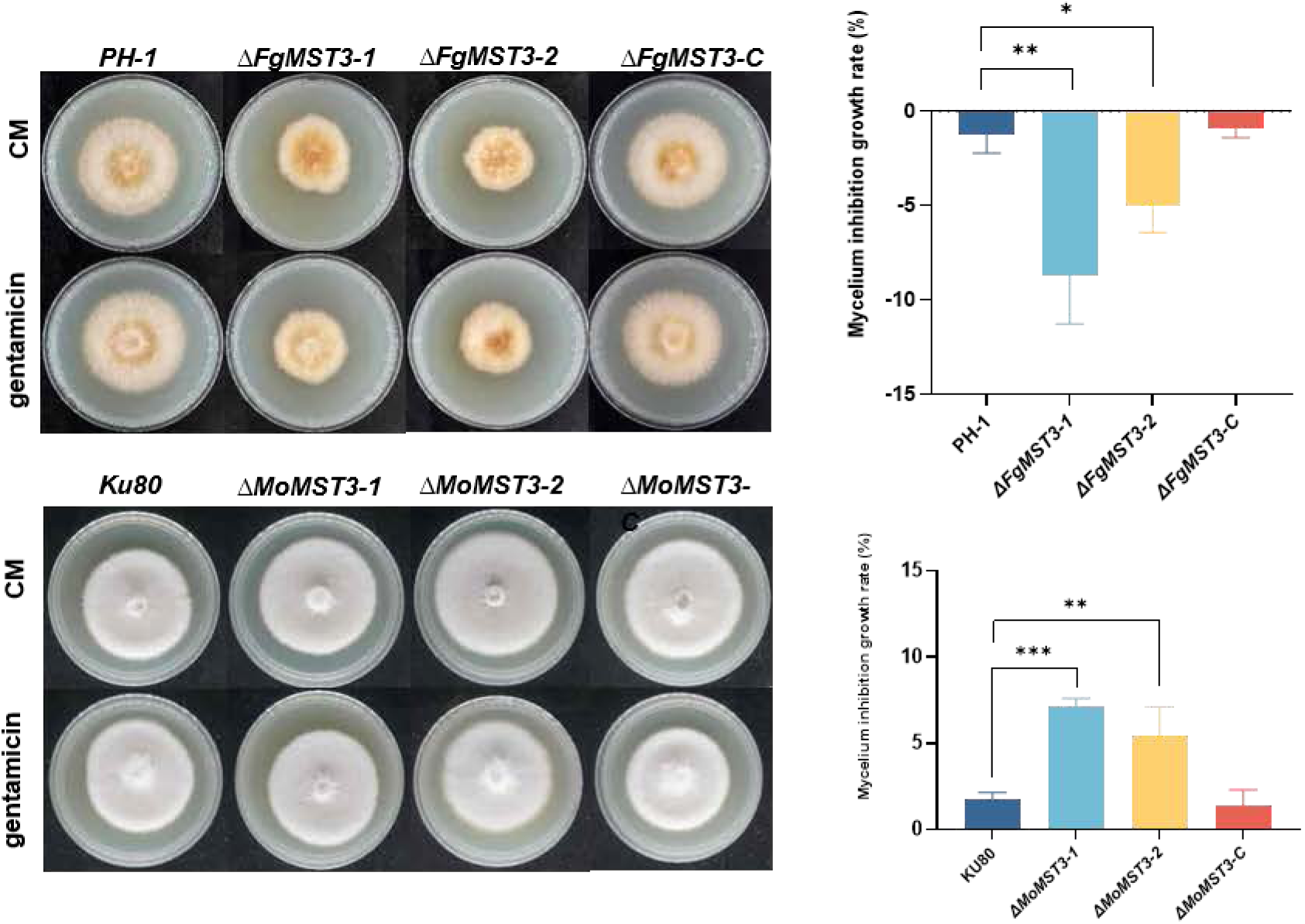
Inhibition of growth by gentamicin^+^ for the different *F. graminearum* and *M. oryzae* strains. Colony morphology of *F. graminearum* strains in the stress medium (A) and inhibition of colony growth on the medium, colony diameter for PH-1 on control medium without stressing agent minus colony diameter on stress medium gives colony growth inhibition (the larger the value, the more inhibited by the stress medium) (B). Colony morphology of *M. oryzae* strains on the different stress media (C) and inhibition of colony growth on the same media, colony diameter for KU80 on control medium without stressing agent minus colony diameter on stress medium gives colony growth inhibition (the larger the value, the more inhibited by the stress medium (D). Error bars indicate SEMs, and *, **, ***, and **** symbolize P_same_<0.05, <0.01, <0.001, and <0.0001, respectively, for the null hypothesis that these measurements are the same as for the respective background strains.

### Effect of the MST3 mutations on fungal innate immunity

A vacuolar iron siderophore transporter, FgMirA, had previously been identified as a potential innate immunity reporter gene strongly and quickly upregulated (1h after bacterial MAMPs exposure) as a potential innate immunity defence effector removing iron from the cytosol and transporting it into the vacuole for storage. Its activity can create a strong sink for the removal of iron from the environment when the fungus is challenged by NSAMPs from surrounding bacteria (5). A fast restriction of bacterial access to iron is a classic part of innate immunity since iron is necessary for many enzymes needed for detoxification of ROS produced by the eucaryotic host, as well as for the host to withstand the ROS produced by its own innate immunity responses to kill invaders (30). We first checked that the vacuolar FgMiraA protein (FGSG_00539) is rapidly upregulated in response to bacterial NSMPs. For that, we constructed a *F. graminearum* reporter strain expressing the FgMirA-GFP protein in the same way as we constructed the FgMst3-GFP complementation strain. We then tested the FgMirA-GFP subcellular localization and regulatory responses when the fungus became exposed to *Esherichia coli* outer membrane vesicles (OMVs) obtained from starving bacteria shedding OMVs carrying a range of NSMPs, imitating the conditions in the soil rhizosphere where *F. graminearum* comes in contact with wheat roots and can cause seedling blight (20). The OMVs are most likely internalized by always present fungal endocytosis (31) for fungal NLRs to become exposed. It was obvious from these experiments with the reporter strain that the strength of the FgMirA-GFP signal responds almost immediately to the *E. coli* NSMPs (**Fig. S8**). Since sucrose, for a fungus, is a potential NSMP (5), we tested the reporter system for a small amount of sucrose (0.1 mg/L), too little to be a significant carbon source in the environment that can potentially serve as a plant NSMP, telling the fungus that it has entered the plant inner rhizosphere or the plant apoplast (5) (**Fig. S9**). These microscopy methods only test a small fungal biomass and are very tricky to perform. We thus tested the transcriptional responses of the *FgMirA* gene to the similar NSMPs relative to a water control using RT-qPCR. As shown in **Fig. S10,** the reporter gene shows clear responses to both NSMPs, although the response to the cocktail of bacterial NSMPs on/in the OMVs was stronger than to the single plant NSMP (sucrose).

Finally we could then test if FgMst3 could be active in fungal innate immunity by testing the *FgMirA* reporter gene transcriptional responses to *E. coli* NSMPs added as OMVs of the *PH-1* strain and compare that with the transcriptional responses to the same treatment of the two *ΔFgMST3* strains (**Fig. 11**). Already after 1 h, the transcription of *FgMirA* was on average 8 times higher in the *PH-1* strain than in the *ΔFgMST3* strains, and that difference rose to 60 times the expression in the *ΔFgMST3* strains when exposed for 2 h.

**Figure 11.**
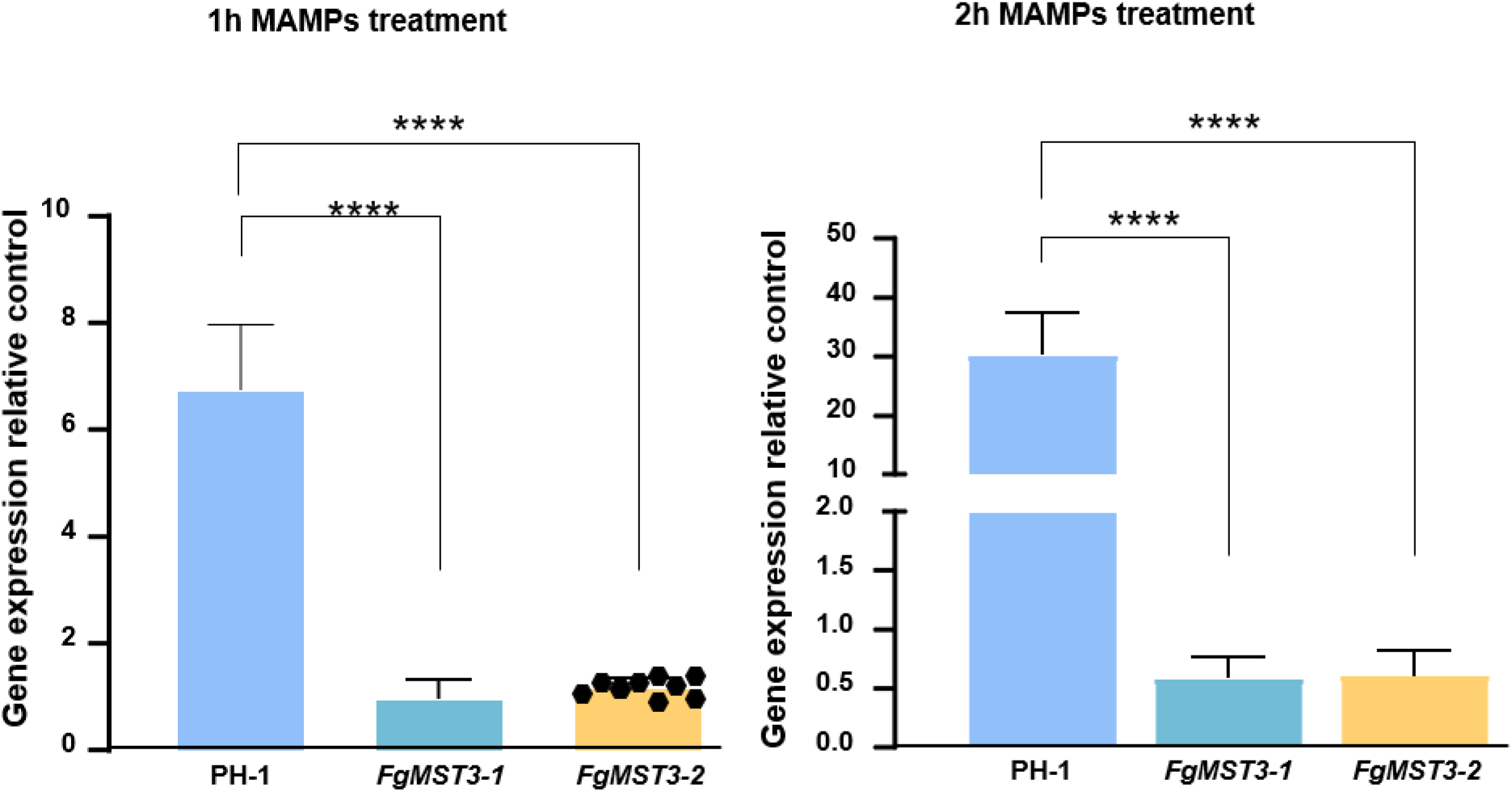
Fast reporter gene expression increases relative water controls for different *F. graminearum* strains after bacterial NSMPs addition. Gene expression after NSMPs treatment for 1h relative to equal water treatment is about 8 times stronger and increases to 60 times stronger after 2 h (B). Error bars indicate SEMs for the 9 replicates marked with black symbols, and *, **, ***, and **** symbolize P_same_<0.05, <0.01, < 0.001, and <0.0001, respectively, for the null hypothesis that these measurements are the same as for the PH-1.

In conclusion, the deletion of the *F. graminearum MST3* gene substantially decreases or even nullifies fungal innate immunity responses to the bacterial OMV treatment, suggesting that FgMst3 proteins relay signals downstream of possible fungal NSMP receptors.

## Discussion

The phylogeny of the different MST3-like protein sequences clusters mammalian sequences together with Dictyostelium and archaea, which could indicate that MST3 proteins are evolutionary as old as innate immunity and developed at the origin of eukaryotic organisms (**Fig. 1**) (7). The yeast ste20 kinases differ somewhat and seem to be a group that has evolved, maybe for different cellular roles. Yeasts are unicellular, and the most studied strains derived from beer fermentation without many bacterial challenges might have lost the most innate immunity, including inflammasome formation and triggering apoptosis to save the mycelium if pathogen invasion is severe in one hyphal compartment. Apoptosis is present but works differently in baker’s yeast and is executed by a metacaspase (32). The highly conserved protein PARP, needed to repair damaged DNA, is rapidly cleaved and inhibited by the apoptosis-activated caspase activities (33), and is not present in yeasts as *Saccharomyces cerevisiae* (34). A PARP was, however, recently found in the environmentally highly genetically competent yeast *Yarrowia lipolytica,* which has a less-reduced number of genes in its genome compared to other ascomycetes, than is the case for *S. cerevisiae* (35). In filamentous fungi, the PARP gene seems more generally present, as well as classic caspase-like protein cleavages by metacaspase (36). Although baker’s yeast is an excellent model for basic metabolism, it seems to fall short of the multicellular (e.g., multinuclear) strategizing seen in filamentous fungi that is important for conidia formation by emptying vegetative mycelium for production of conidia and to spread to a more nutritious environment (34, 36). Filamentous fungi are also known to have NLR like proteisn of types that should be able to sense bacterial and plant NSMPs. Some of these are known as Het proteins involved in heterogenous incompatibility proteins triggering local cell death in the model fungus *Podospora anserina* (14–17). Similarly, and different from yeast fungal thalli (the multinucleate mycelium), can “migrate” from a hostile volume in the soil or search for a more “rewarding” volume in the soil by recycling nutrients to support the migration through “barren” soil volumes (37–40).

In light of these capacities of filamentous fungi with a general high competence to thrive and grow in competition with bacteria interacting with hyphae, both pathogenic and beneficial to the fungus (41), filamentous fungi should need signaling proteins like the well conserved Mst3 kinase to relay the signals from NLR-receptors to innate immunity effectors like siderophore transporters to remove iron from their surroundings (5). The lack of responses to bacterial NSMPs for the *ΔFgMST3* strains compared to the PH-1 background strain is in line with this and indicates that there must exist, so far non-identified, NLR proteins being exposed to endocytosed bacterial NSMPs, recognizing them and triggering downstream responses, as the quick upregulation of the FgMirA (**Fig. 11**) shows. Candidates for such NLRs can most probably be found among the approximately 100 identified *F. graminearum* STAND proteins (17) that are implicated as possible bacterial NSMP receptors recognizing bacteria (15, 16).

Bsr1/Irak4 kinases are short, while the MST3 kinases are much longer, and their C-terminal part behind the N-terminal kinase domain contains an NLS sequence and a NES sequence, and between them a cleavage site for caspase that, when cleaved, further activates the kinase (25). It was thus expected that the two fungal MST3-like kinases should have a domain structure like the human MST3-kinase if they have a similar and conserved function in innate immunity (**Fig. 2, S1, and Supplementary file SF1**). On the other hand, this similarity should not necessarily apply to *S. cerevisiae* since it has a very reduced genome and typically lacks genes for proteins important for multicellular organisms and for existing in multiple natural environmental settings (35). Our results suggest that MoMST3 and FgMST3 could have a function similar to the mammalian MST3 gene and relay signals from NLRs downstream to innate immunity, as the deletion of the *FgMST3* gene severely affected the innate immunity effector reporter gene (*FgMirA*) transcriptional responses (**Fig.11**).

*MST3* gene deletions in both plant pathogenic fungi had negative effects on pathogenicity, most probably since the deletions affected conidia production with fewer conidia, changed the dynamics of conidia germination, and reduced colonization of the plant host tissues (**Fig. 7-8**). Interestingly, both the MST3 mutants germinated faster than their respective background strains (**Fig. 5**). We had deliberately selected two fungal pathogens with different natural bacterial environments. *F. graminearum* has wet conidia spread by raindrop splashing and meets bacteria in the soil and rhizosphere/rhizoplane of plants, while *M. oryzae* mainly has dry melanized conidia and meets bacteria in the sunlit phylloplane of grass leaves. Because of these differences, the effect of MST3 deletions on genes downstream of *MST3* is expected to be different (**Fig. 6, 9, 10**). Since *F. graminearum* lives in soil when not infecting a plant, it should be more dependent on its capacity to withstand biotic stresses imposed by bacteria in the soil and rhizosphere. *M. oryzae* should, on the other hand, be more dependent on a properly formed melanized cell wall and ROS defences since the environment on the leaves is very oxidative and prone to high ROS levels, also caused by intense sunlight. Inside the leaves, conditions are better controlled as long as there are nutrients available to run ROS defences. The largest difference we found in the change of sensitivity for the *ΔMST3* strains in comparison with the background strains was in Zn^2+^ resistance, where the *ΔFgMST3* strains were more tolerant to Zn^2+^, although zinc can work as a fungicide towards *F. graminearum* (42), and they were also very tolerant to the antibiotic gentamicin. *ΔMoMST3* strains were less tolerant to both treatments compared to their respective background strains (**Fig. 9-10**). The gentamicin treatments even stimulated the growth of *F. graminearum,* and the *ΔFgMST3* strains were even more stimulated than the PH-1 strain. At present, we have no plausible explanation for that stimulation.

## Conclusion

The detected reactions of the two tested fungi to Gram-negative bacterial NSMPs common in plant rhizospheres and the disappearance of these reactions when the *MST3* genes were deleted indicate that the encoded fungal Mst3 protein relays signals downstream from hitherto unknown fungal NLR-receptors of types postulated to be active in recognizing invading or interacting organisms, triggering fungal innate immunity. The two identified *MST3* genes seem to have similar roles in both fungi and play a role in the normal plant infection processes of both these pathogens. The negative effect on pathogenicity of deleting the *MST3* genes in both fungi might be caused by a lack of recognition of unknown plant NSMPs, which normally prepare the fungus for a successive plant invasion where it meets plant immunity responses. Future work should try to identify NLR receptors as well as plant NSMPs to better understand how fungal non-self-recognition works and if it can be inhibited or manipulated, as well as teach us something about human innate immunity, since the fungal version seems more similar to human innate immunity than plant innate immunity (5).

## Materials and Methods

### Websites and programs that were used for the initial bioinformatics work

We used NCBI for the download of protein sequences and for BLAST similarity searches, as well as conserved domain descriptions. Multiple alignment of homologs was performed and visualized using the NCBI COBALT multiple alignment tool at the NCBI website. For phylogeny inferences of the found protein sequences, we used NGPhylogeny.fr (43) (https://ngphylogeny.fr/) using the ‘One click’ mode for tree building. ScreenCap 3 (https://scap.cbrc.pj.aist.go.jp/ScreenCap3/index.php). was used to predict putative caspase 3 cleavage sites. Finally, the protein conserved domain structures were visualized using the free tool for Protein domain structure visualization: DOG 2.0 (44), available for download at http://dog.biocuckoo.org/. The detailed step-by-step procedure of our bioinformatics work is available (**Supplemental File SF1)**.

### Fungal strains, plant varieties, and plasmids

The fungi we used for the study are both well-studied model plant pathogenic Ascomycetes. We used *Fusarium graminearum* (teleomorph *Gibberella zea*) wildtype strain PH-1, and *Magnaporthe oryzae* (formerly *Magnaporthe grisea*) (Teleomorph *Pyricularia oryzae* wildtype strain of Guy11. The Guy11 strain we used has the gene *MgKU80* required for non-homologous end joining inactivated, strain Ku80, to aid gene targeting (45). Information about genes and proteins for both strains can be found at NCBI (*F. graminearum* NCBI:txid229533, *M. oryzae* NCBI:txid242507). Plants used were wheat (*Triticum aestivum*) variety Jimai 22, rice (*Oryzae sativa*) variety Co39, and barley (*Hordeum vulgare* subsp. *vulgare*) variety Xiyin No. 2. The plasmid used for knockout was PCMB containing Ampicillin resistance and HPH (Hygromycin resistance) for selection of putative successful knockouts. For complementations, we used plasmid PKNT-GFP containing Ampicillin resistance and G418 resistance to aid in the selection of putative successful complementations.

### Gene knockout mutations and complementation of knockout strains with *MST3-GFP* constructs

Targeted gene deletion and complementation were performed as described (Chen et al., 2023). For making respective *ΔFgMST3* and *ΔMoMST3*, ∼800 bp 5’ and 3’ flanking regions were amplified and fused to a hygromycin resistance cassette (HYG), followed by polyethylene glycol (PEG)-mediated protoplast transformation. Primary transformants were selected on hygromycin (200 μg/mL) and validated by Southern blot. Complementation constructs (1.5 kb native promoter + GFP + FgMST3 or MoMST3) were cloned into pKNT and reintroduced into *ΔFgMST3* or *ΔMoMST3*. GFP fluorescence was confirmed via confocal microscopy (Nikon A1, Japan).

Successful strain constructions were further verified through PCR amplifying parts of flanking regions + knockout inserts and getting predicted band sizes, in combination with our own Gong/Olsson qPCR technique (GO-qPCR for short) to determine the *HYG/Tubulin* gene ratio in the mutants to detect possible ectopic integrations of the *HYG* gene used as a selection marker, using standard primers we have in the lab for these genes, amplifying only inside their exons. GO-qPCR developed for this study is a faster and cheaper alternative to Southern blotting, which also quantifies the number of ectopic integrations in the genome. Using this method makes it less advantageous to use the KU80 strain to avoid multiple ectopic integrations through NHEJ, since screening for multiple integrations could simply be done as a first measure after transformation. This is feasible since only small amounts of gDNA are needed for qPCR, and could be done when colonies have first been obtained on the selective medium, using small samples taken directly from parts of colonies appearing on the hygromycin plates. Thus, subculturing/isolation needs only to be done from colonies showing one *HYG* gene integration in their genomes.

### **G**rowth of strains and stress treatments

*F. graminearum* and *M. oryzae* mycelial disks were used as inoculum for CM, CMII, SYM, and MM agar plates that were incubated at 26 °C for 3 days (*F. graminearum*) or 7 days (*M. oryzae*) before the plates were photographed, and the colony diameters were measured, and relative inhibition rates were calculated (46). For in vitro stress sensitivity assays, fungal strains were inoculated on CM solid media with one of the following additions 0.02% w/v calcofluor white (CFW), 1M NaCl, 1M KCl, 1mg/mL Congo Red (CR), 0.01% (w/v) sodium dodecyl sulfate (SDS), 0.03% w/v H2O2, or 1 M Sorbitol. Mycelial disks were used as an inoculum for the stress treatment plates and CM control plates, and they were incubated, photographed, and measured as before for the two pathogens, respectively.

### Conidia production and germination measurements

Strains of *F. graminearum* grown on CM agar media were used as inoculum to shake cultures (180 rpm) in CMC liquid medium and incubated at 25 °C. Conidia were counted after 3, 6, and 9 days of incubation using a hemocytometer. Disks taken from CMII cultures of *M. graminearum* strains were used to inoculate rice bran plates and incubated inverted for 7 days before the aerial mycelium was scraped off into a clean disk and incubated at 26 °C under near ultraviolet illumination to promote conidia formation for 2 days. After incubation, the conidia were washed off and counted using a hemocytometer.

For germination experiments, 10 mL conidia suspension 2*10^4^ /mL was pipetted to a hydrophobic microscope slide and incubated in a high-humidity chamber in the dark at 26 °C, and conidia germination and appressorium formation (*M. oryzae*), were investigated by microscopy after 2, 4, 6, and 8 hours (28).

### Pathogenicity testing of *F. graminearum* strains

**Wheat coleoptiles** grown from alcohol sterilized seeds (cv. Jimai 22), on sterilized filter paper wetted with sterilized water, were used for the testing. A *F. graminearum* conidia suspension containing 2 × 10^4^ conidia/mL from a liquid CMC medium incubated for 3-4 days was used as inoculum. The tip of the coleoptiles was cut and inoculated with 10 mL of the conidia suspension, and the coleoptiles were placed in a tray and covered with a plastic wrap. These trays with inoculated coleoptiles were then incubated at 26 °C with an 8/16 dark/light period for 3 days before lesion sizes were recorded.

**Wheat heads** of field-grown winter wheat plants grown from seeds that were sown in mid-December were inoculated at the flowering stage in April-May the following year, and were used for the testing. Even-sized small disks cut from colonies of the *F. graminearum* strains were used to inoculate the wheat heads. One fungal disk was placed on the 4^th^ or 5^th^ floret from the bottom of each wheat head. Each head was covered with a plastic bag for 3 days and then removed. Then, after about 2 weeks, the wheat heads were cut off, photographed, and the levels of wheat head infection were estimated.

### Pathogenicity testing of *M. oryzae* strains

Barley leaves (cv. Xiyin No. 2) with similar growth characteristics were selected from 6-7-day-old plants, and placed in a moist chamber made of a 15 cm Petri dish. A spore solution containing 2 × 10^4^ conidia/mL was added to the barley leaves in spots of 15 mL/spot. The moist chamber containing the inoculated leaves was then incubated at 26 °C in the dark for 24 h, followed by 12/12 dark/light periods for 3-5 days before lesion sizes were recorded.

Rice leaves from 4-week-old rice seedlings (cv. CO39) were spray-inoculated with 3 × 10^4^ conidia/mL. Inoculated plants were maintained at 26°C (>90% humidity) for 5-7 days. Disease severity was scored on a 5-grade scale based on lesion size: grade 1 (<0.05 mm), grade 2 (0.05-0.2 mm), grade 3 (0.2-0.3 mm), grade 4 (0.3-0.4 mm), and grade 5 (>0.4 mm) (47).

Rice plants (cv. CO39) were grown until the 3-leaf-1-heart stage, and leaves to be tested were excised for press-injured inoculation (48). In short, 2 mm diameter press-injured spots were made using a pressing machine (Fujihara Co.), and 10 mL of a conidia suspension containing 5 × 10^5^ per mL was added to each injured spot, and the spots were sealed with transparent tape. The leaves were then incubated at 26 °C with a 12/12 dark/light period for 10 days before lesion sizes were recorded.

### Confocal microscopy observations of the subcellular localization of Mst3 proteins in the complement strains expressing the FgMst3-GFP or MoMst3-GFP proteins

Subcellular localization was analyzed using a Nikon A1 confocal microscope. EGFP-tagged strains were cultured on CM II, fixed in 4% paraformaldehyde, and imaged using blue light excitation (EGFP) (Argon laser 488. Excitation filter:470/40, Dichromatic mirror: 495LP, Barrier filter: 515/30 (the light imaged).

### Obtaining. *E. coli* OMVs containing bacterial NSMPs

OMVs from overnight cultures of *E.coli* (strain DH5a) grown in 5 ml Luria Broth LB at 37 °C were centrifuge-harvested at 10000 rpm, washed twice in sterile double distilled (DD) water, and finally resuspended in 1ml DD water. Then the cells were incubated for 1hour at 37 °C to release OMVs that are typically formed during starvation. The OMVs were finally harvested from the supernatant by centrifugation at 5000rpm to remove most cells. The supernatant with OMVs was sterile filtered through a 0.3 μm filter. The resulting sterile OMV suspension can be stored in the fridge at 4 °C until use. For treatments for microscopy, the OMV suspension was diluted approximately 10 times, and the washed OMV suspension was added to washed and starved hyphae and incubated on the microscope slide for 1 or 2 test hours before the cover slip was put on and the slide was immediately investigated in the confocal microscope.

For RT-qPCR, the OMV concentration used to expose the fungus was diluted 10 times before the starved hyphae were exposed, and the fungal mycelium was incubated for 1 or 2 h before immediate harvest of mRNA for RT-qPCR assessments. The procedure was as follows: The fungal hyphae to be treated with OMVs were produced in well-aerated liquid DFM medium cultures incubated for 3-4 days at 26 °C. After that incubation, the remaining medium and metabolites in the medium were removed by filtration, and the hyphae on the filter were transferred by shaking the filter in N-deficient DFM overnight. The mycelium was filter-washed again with DD water and finally transferred to a 50ml centrifuge tube containing 1mLsterile DD water, and 1 mL OMV suspension was added, exposing the fungus to OMVs containing bacterial NSMPs. The exposure of the starved fungal hyphae to OMVs from starved bacteria in DD-water mimics the starving conditions in the soil rhizosphere, where the fungus meets bacteria, takes up the few nutrients that are available, and interacts with the biotic and abiotic environment also through endocytosis.

## Acknowledgements

Dr. Mengtian Pei is thanked for her help in guiding Shanshan Gong in her laboratory work. This work has been supported by funds from The Key program of Science and Technology in Fujian province, China (2024NZ029027), the Fujian Provincial Natural Science Foundation (2026J001341), and the Fujian Agriculture and Forestry University Science and Technology Innovation Special Fund (KFB23018A).

## Author contributions

SG First author, made MST3 mutants and experiments, including OMV extraction, and manuscript writing.

XL, co-first author, made the reporter strain, tested the technique to extract OMVs, and tested whether RT-qPCR works with the reporter gene.

SL refined the RT-qPCR technique for measuring reporter gene responses to OMVs so that it works reliably.

SO Co First author and co-corresponding author, research idea, identification of reporter gene, OMV purification technique development, bioinformatics, manuscript writing, Supervision

GL Co-supervision ZW Co-supervision

YL, Corresponding author, provided funds and Co-Supervision.

## Supplementary figs

**Figure S1.**
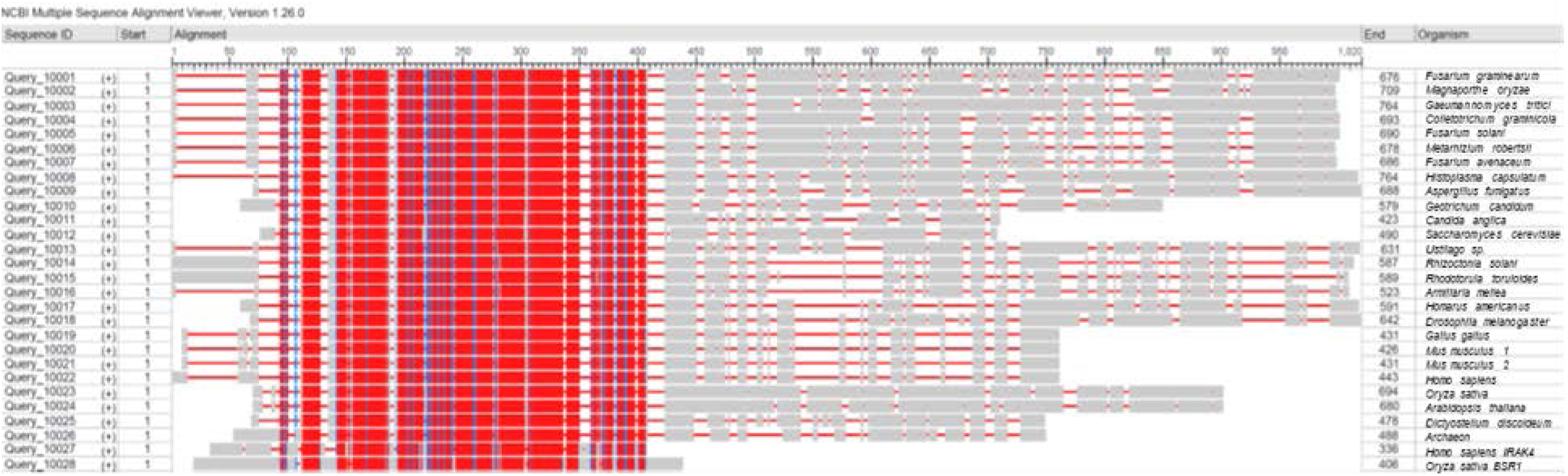
Alignment of the MST3-like proteins in Figure 1. In the alignment, the kinase part of the protein aligns well for all proteins, although the longer C-terminal part is absent from the IRAK4 and BSR1 proteins known to be involved in relaying signals from animal and plant TLR receptors.

**Figure S2.**
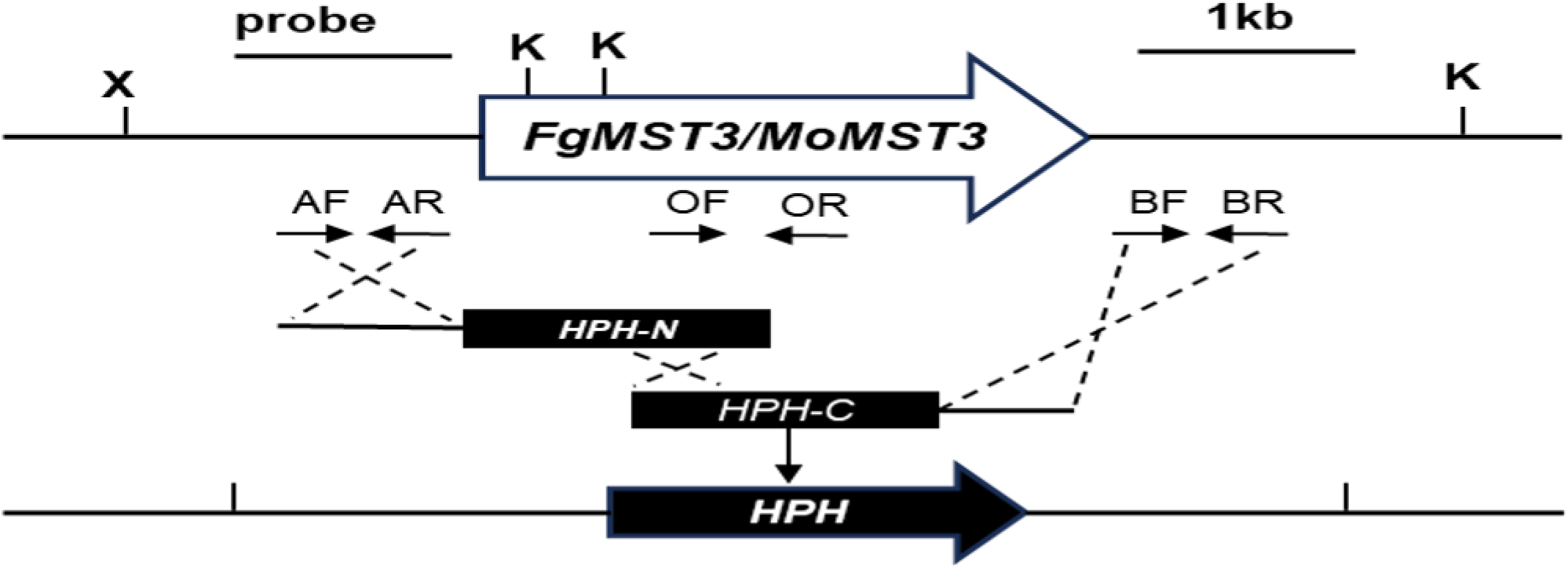
**Deletion strategy for *FgMST3* and *MoMST3***

**Fig S3.**
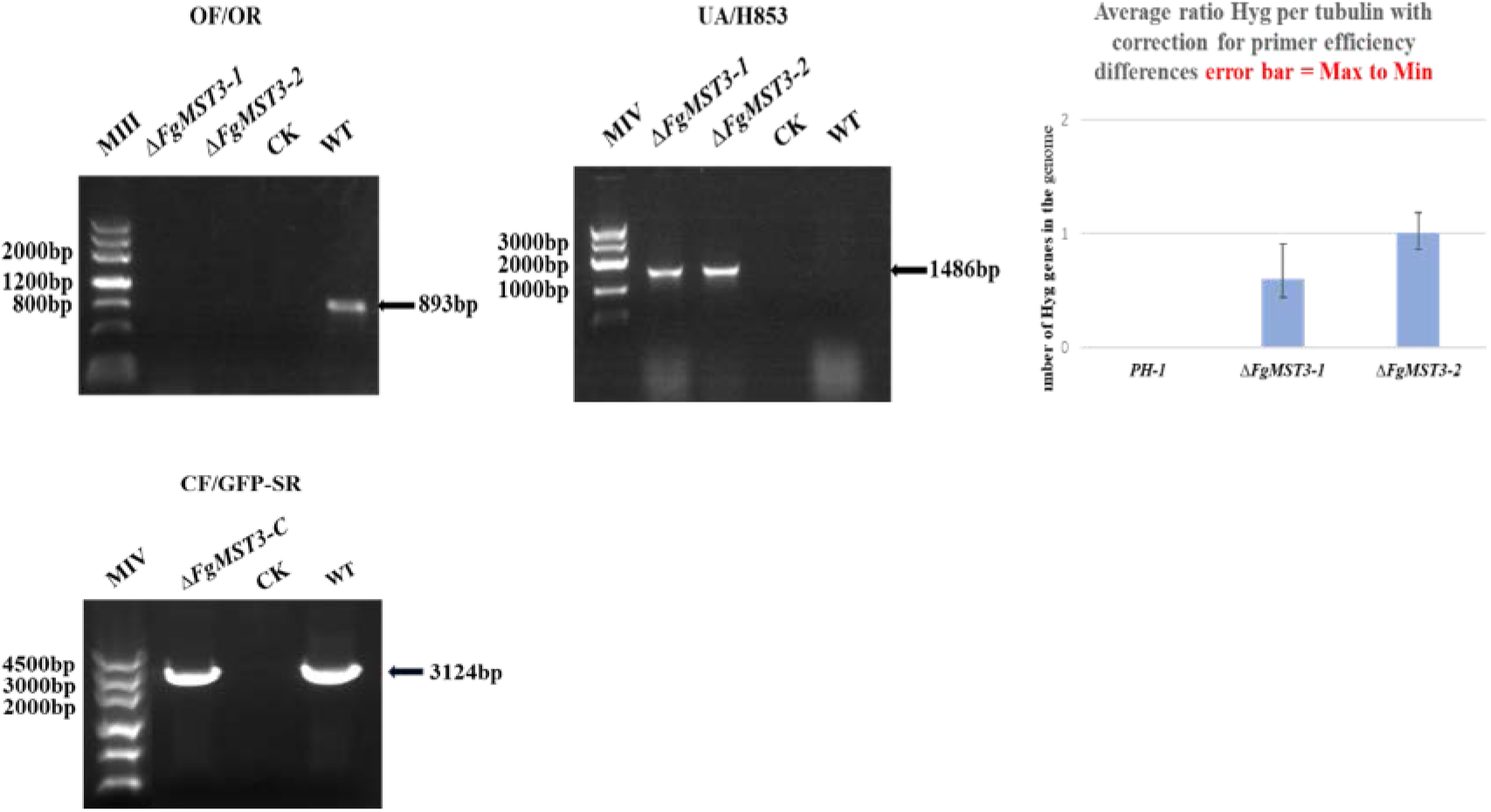
Confirmation that the constructed strains have the right new genes in their genomes and that there are no ectopic insertions of the gene used as a selection marker.

**Figure S4.**
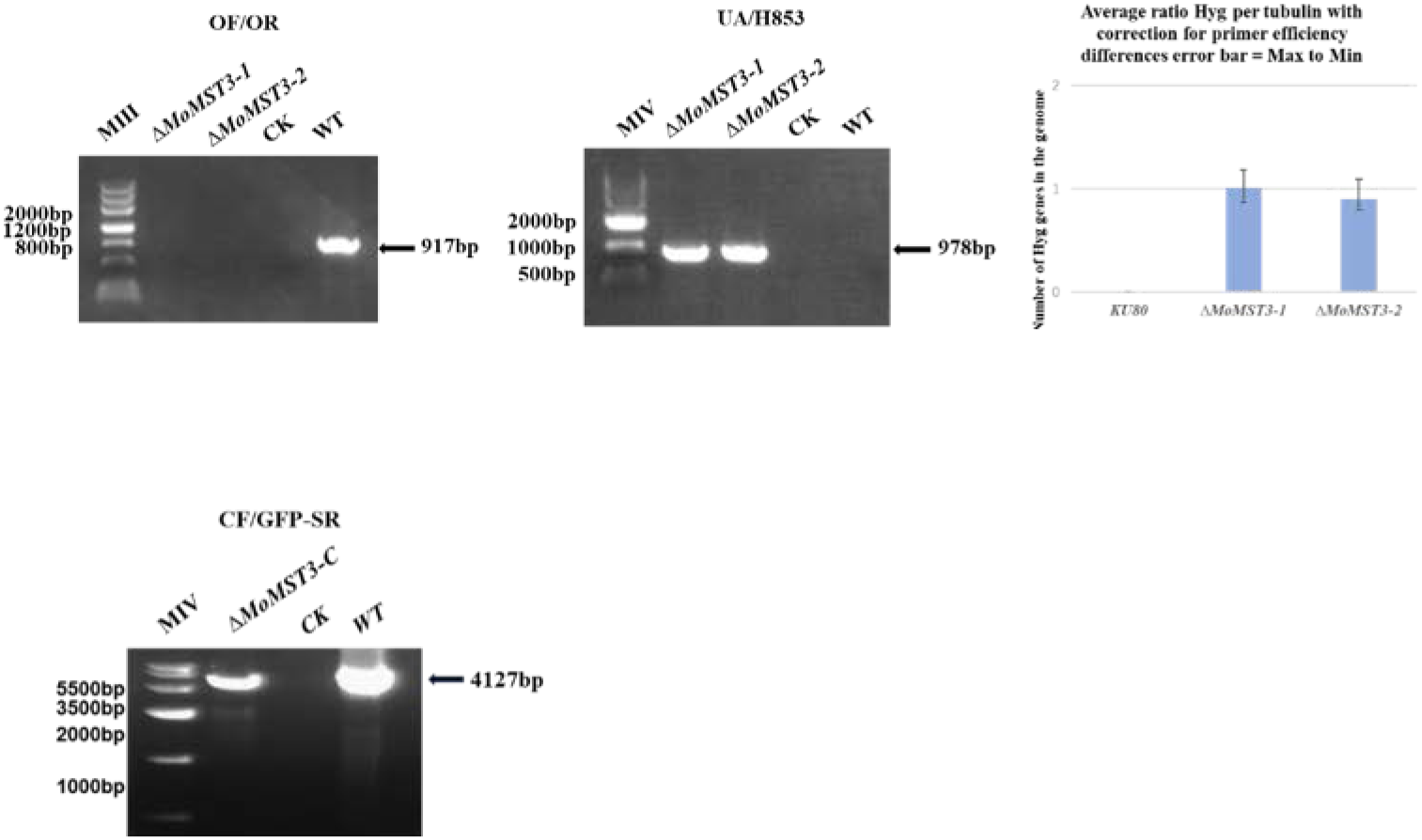
Confirmation that the constructed strains have the right new genes in their genomes and that there are no ectopic insertions of the gene used as a selection marker.

**Figure S5.**
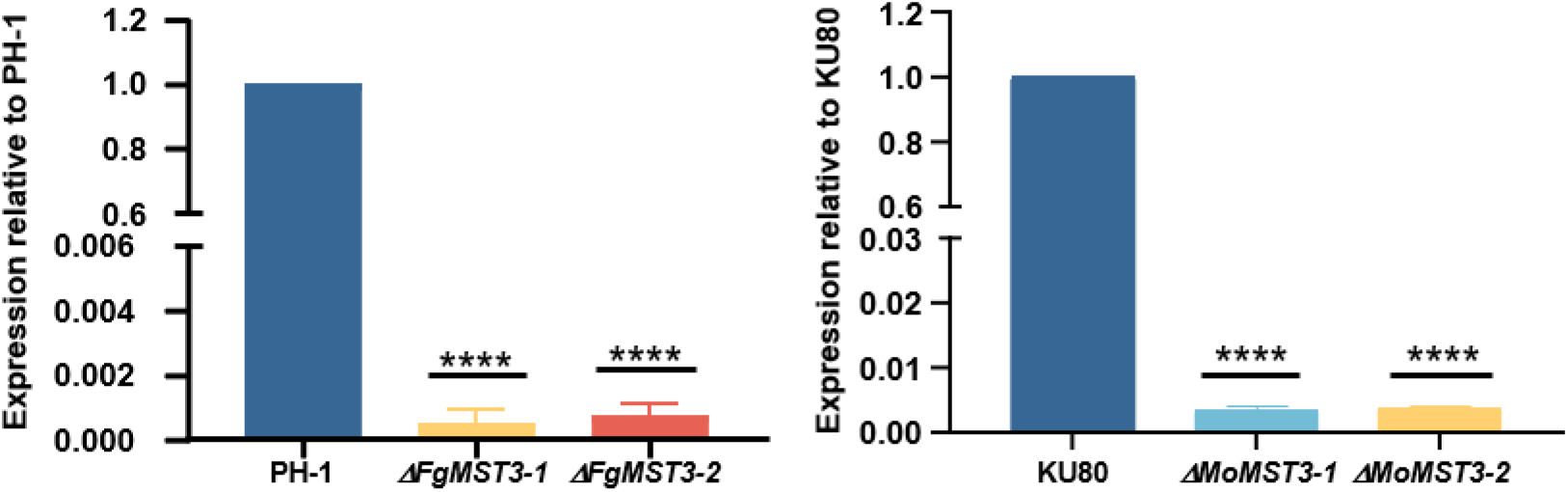
RT-qPCR confirmation that there is no residual expression of the deleted genes in any of the mutants. Error bars indicate SEMs and *, **, ***, and **** symbolize P<u>sanv^</u>O OS, <0.01, <0.001 and <0.0001, respectively, for the null hypothesis that these measurements are the same as for the PH-1 or KU80, respectively.

**Figure S6.**
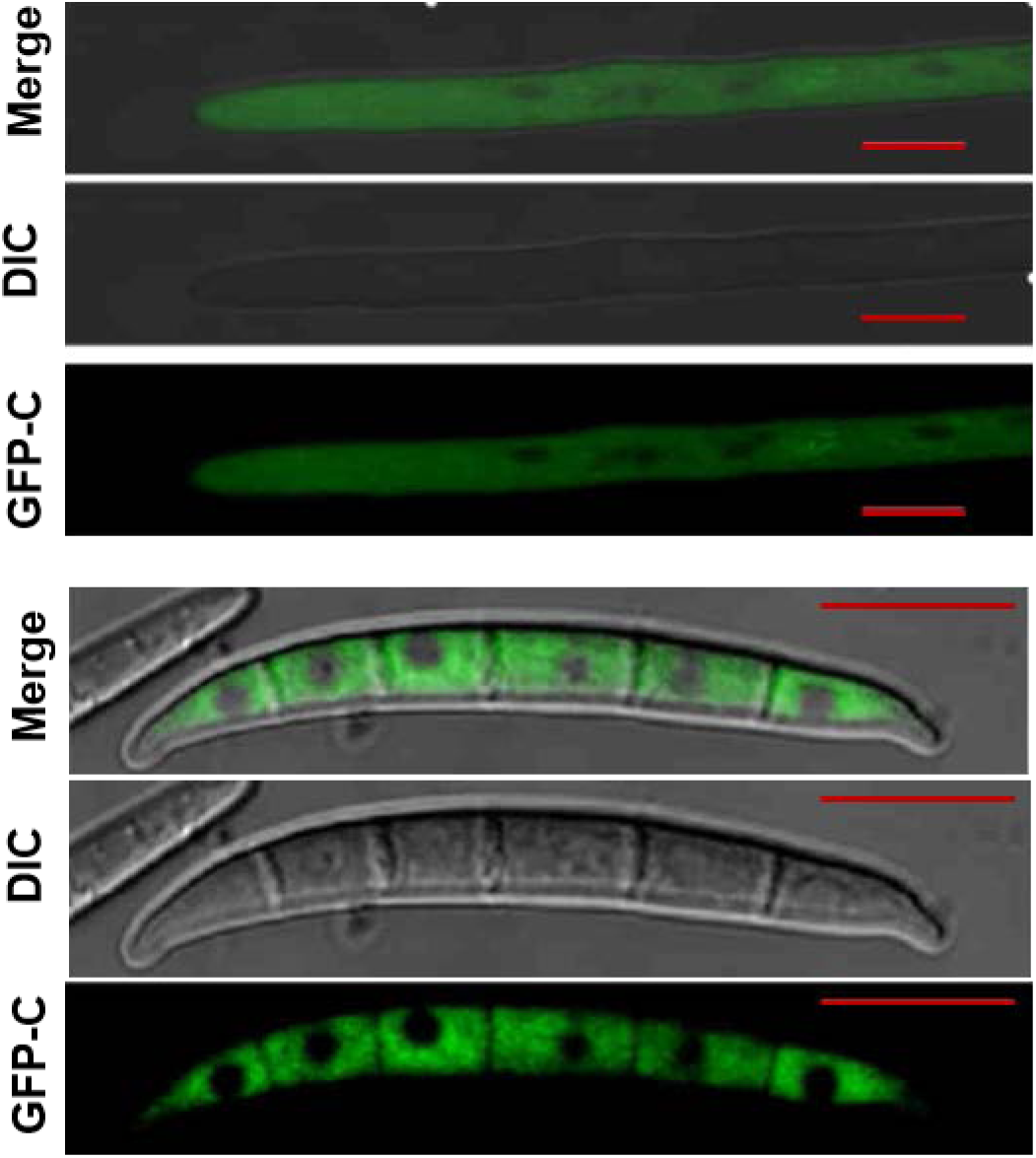
Subcellular localization. FgMst3-GFP localizes to the cytoplasm in hyphae (A) and conidia (B). Size bars = 5 pm (spore images), and 10 pm (hyphal images)

**Figure S7.**
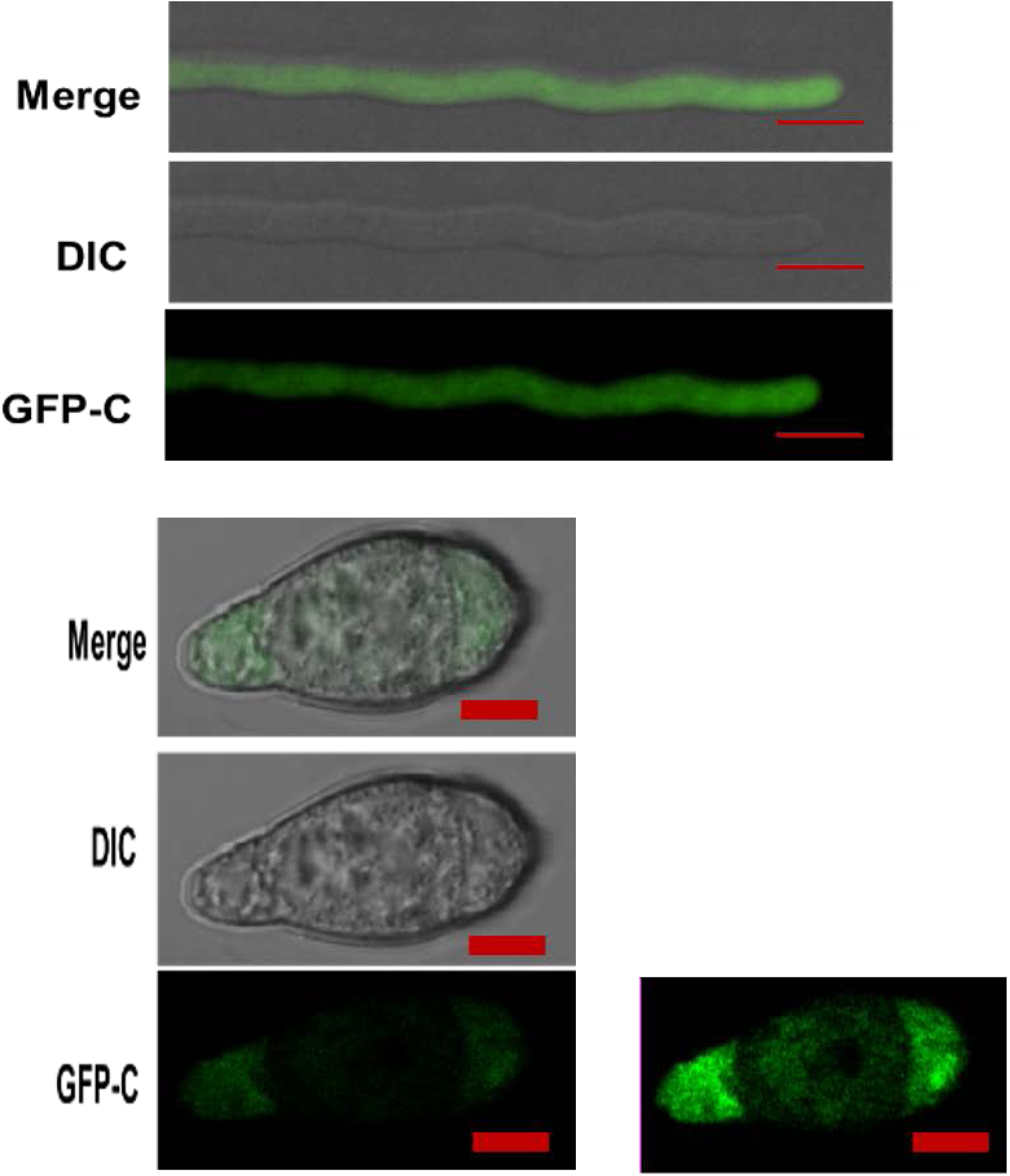
Subcellular localization. MoMst3-GFP localizes to the cytoplasm in hyphae (A) and conidia (B). The central compartment of the conidium is heavily melanized and blocks the blue light needed to excite GFP but the signal appear clearly cytoplasmic as seen in the more enhanced image. Size bars = 5 pm (spore images), and 20 pm (hyphal images)

**Figure S8.**
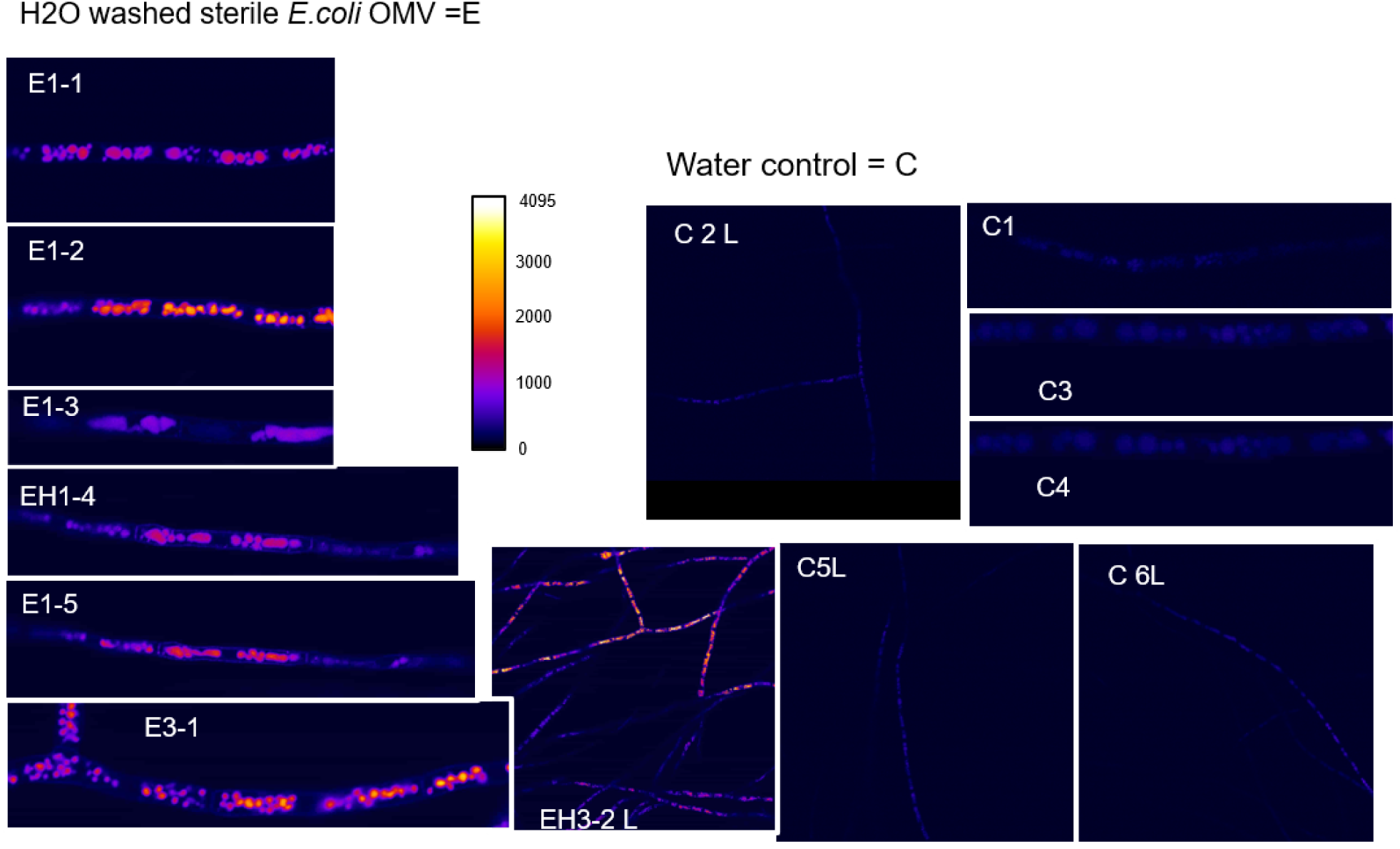
**Response of the reporter strain FgMirA-GFP when treated with NSMPs from *E. coli.*** The reporter strain was washed and starved in sterile water before treatment. Treatments: H_2_O: washed sterile *E.coli* OMV suspension treatments for lh, 5 replicate treatments El-5. H_2_O-washed sterile *E.coli* OMV suspension treatments for 3h, 2 replicate treatments E31-2, seen in low magnification L. H_2_O control: H_2_O for 1 to 6 hours C1-C6. C2, C5 and C6 seen in in low magnification L. NOTE: The lookup table Fire was used to better visualize the large differences of the responses to human eyes. The scale in the center shows how the Fire lookup table represents the 12-bit confocal images with 4095 light levels.

**Figure S9.**
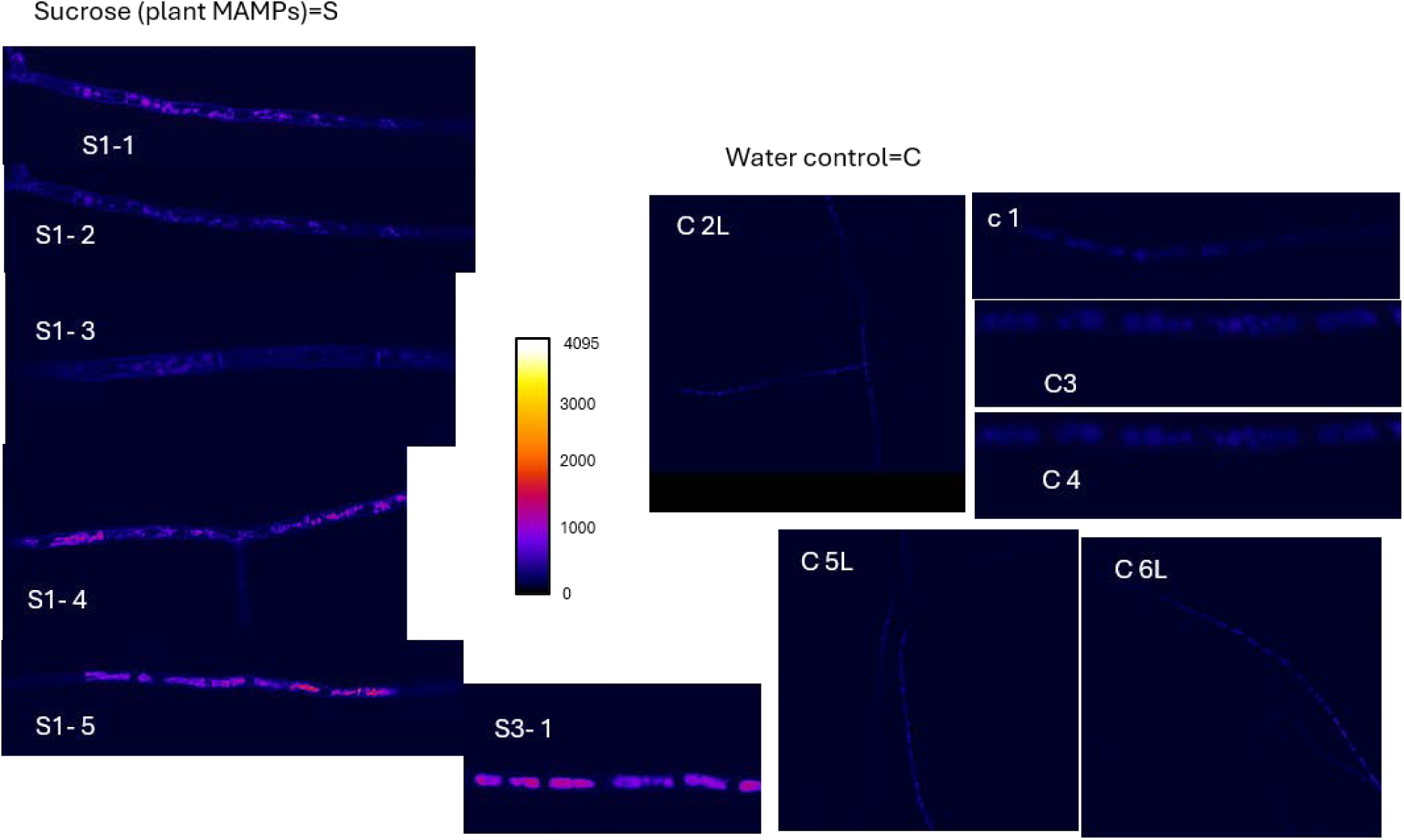
Response of the reporter strain FgMirA-GFP by when treated with a plant NSMP. The reporter strain was washed and starved in sterile water before treatment. Treatments: Sterile sucrose solution (0.1 mg/L) treatments for lh, 5 replicate treatments Sl-5. Sterile sucrose solution (0.1 mg/L) treatments for 3h. Control: H_2_O for 1 to 6 hours C1-C6. C2, C5, and C6 seen in low magnification L. NOTE: The lookup table Fire was used to better visualize the large differences of the responses to human eyes. The scale m the center shows how the Fire lookup table represents the 12-bit confocal images with 4095 light levels. As a reference: The apoplast of plants contains about 0.1-5 mM =0.1*360/1000 =0.036-0.18% W/V = 0.36-1.8g/L=360-1800mg/L. Thus O.lmg/L is probably too low a concentration of sucrose for the fungus to grow on but could work as a plant NSMP signal to detect the presence of a plant when the fungus is on the rhizoplane, since sucrose is virtually absent from soil due to its fast consumption by most soil bacteria.

**Figure S10.**
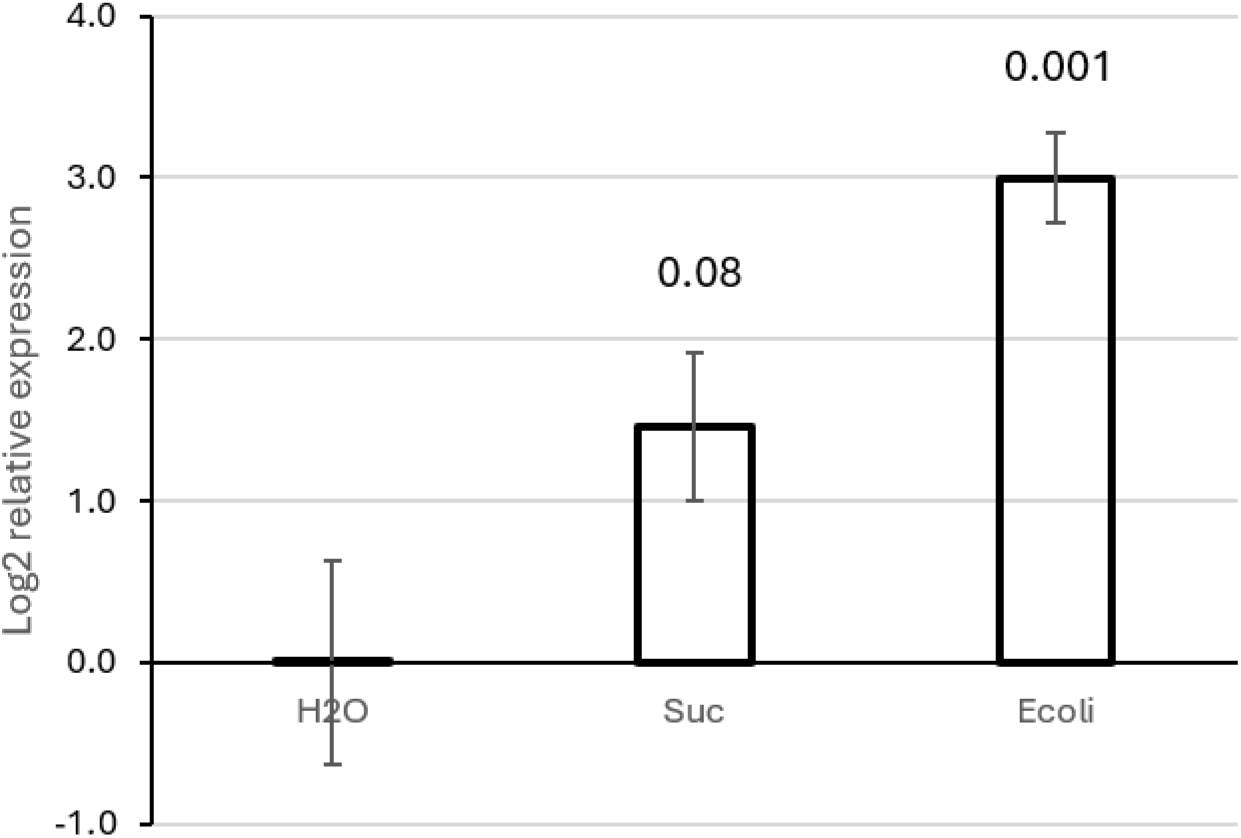
RT-qPCR transcription test if the reporter gene *FgMirA* (FGSG 00539) upregulates in response to plant and bacterial NSMPs. The responses as Log2 gene transcription responses relative/water control was used as the relative measurements. The responses were recorded after lh treatments. H2O = Autoclaved DD water sterile filtered as control treatment. Sue = Hyphae grown in liquid medium, washed and then starved in water was exposed to 0.1 mg/L sucrose. Ecoli = *E. coli* OMVs, or just water as in Fig. S9. The numbers above the bars are the probability for the null hypothesis (P_same_) that the treatments are not different from the control.

