## Supplemental FIle 1 for "Fungal Mst3-proteins are involved in fungal innate immunity needed for the recognition of bacteria surrounding the hyphae, as well as for plant pathogenicity"

### MST3 Supplementary File SF1

Caspase cleavage site prediction at https://scap.cbrc.pj.aist.go.jp/ScreenCap3/index.php

and nuclear localization signal by NLS prediction by comparing the NLS signal in human by aligning the N-terminal part before the first predicted good caspase cleavage prediction (see fig next)

Caspase is predicted to cleave between the Yellow and Red marked AAs. Green marked AAs are the predicted NLS based on the NLS in the human protein, leaving the bold marked N-terminal part containing the kinase domain with the NLS signal that then translocates to the nucleus

>XP_011324316.1 serine/threonine-protein kinase 24 [Fusarium graminearum PH-1] FGSG_05734 MST3 42/4; 3
**MADREHDIEEGNEALDPELLYSKEYCIGGGSFGKVYKGVDKRTGQSVAIKVIDIESAEDEVEDIIQEIAI**
**LSELQSPYVTKYYGSYAKGAELWIVMEFCSGGSCADLMKPGLISEDYIAIIVRELLMGLDYLHTDKKLHR**
**DVKAANVLLSSNGQVKLADFGVSGQLSATMTKKNTFVGTPFWMAPEVIKQSGYDHKADIWSLGITALELA**
**NGEPPYADIHPMKVLFLIPKNPPPRLEGNFTKAFKDFIESCLQRDPKDRPTAKDLLRHPFIRRAKRTTYL**
**TELIERHSRWAAAHKGEDDDNWESVNDGRPPAEHERVDEDMWD**FGTVRLVGDRGGIVNRPGLNILDENAT
NARASRPLESEEDYGEQRREASPMKRDFALQPLETVKAPNSRQSSPQRKAVPQSQYQPPLSPTRALPQTP
LKLPARDSADTPRPLRSVHAPVPESPEYDRELQNELQRDIGMLNLGYDPKESRSPQPQPSQRPPAPPAHL
TRPTTTKQMSLPEIPPYRPSQAVSQQRLPSLQHASTPQVLVPLPGGAQGPRPLPSKNSPSPSDFTAPSAF
PTPSPANPNGELDALNDVIFPALEEALKRRQINLQQVYKPGQNPAQIPPQQQRAEAAHEKLRKLVYKLAH
VCKEIDHYDKAEPVGMGREVGSFLEGLLEEILVRVEPLDEDEDVQS

Probability of caspase cleavage: 0.930483 Site:ARDSAD|T (D430)

Probability of caspase cleavage: 0.849901 Site:DEDMWD|F (D323)

>XP_003719239.1 STE/STE20/YSK protein kinase [Pyricularia oryzae 70-15] MGG_08746
**MGDRDRDYEEGSPDPESLYTKEYCIGGGSFGKVYKGVDKRTGHAVAIKIIDIESAEDEVEDIIQEIAILS**
**ELQSPYVTKYYGSYAKGAELWIVMEFCSGGSCADLMKPGLIGEEYIAIVVRELLLGLDYLHADKKLHRDV**
**KAANVLLAANGQVKLADFGVSGQLSATMTKKNTFVGTPFWMAPEVIKQSGYDHKADIWSLGITALELANG**
**EPPYADIHPMKVLFLIPKNPPPRLEGNFTKAFKDFIELCLQRDPKDRPSARELLRHPFVRHAKKTSYLTE**
**LIERHSRWSSTHKSENDEELD**SQSDDEQQRQQQAANRAAVNEDMWDFGTVRLVGERGHVVHRPGLLNPLD
EAATNARATRSHETEFEQNRQRYRSESPTKRLSQQQLDNDTVKSRQVSPQRRPVPSSPGKTTVQKIEREP
ETPRAVLKPAPISFDSAPSPDYDRELQKQLQREMDGMNIGDMIDPGHKDRTGLIPKTGPASSQAQKPIAH
LGPIKLPEIPPFRGKHAGSSQPAAQPQTRQYQPPQPRQMQQQELPPHPPYRARDDTPEPPQHQQLPTPRP
LQTQALQYQQQQQQQPGTPQHQDFPSPAPAPPNGEMDALNDVIFPALEEALKRRQIRLQQQIQAMPPGAA
SQFAGGAPAPTPRQQRAEAAHEKIRKLVYRVAHMCKEIDHYDKSEPVHMDTGKDAGGTFLEGFLEEIMLR
VEPFDEGEG

Probability of caspase cleavage: 0.849635 Site:NDEELD|S (D301)

Probability of caspase cleavage: 0.796064 Site:NEDMWD|F (D326)

>NP_003567.2 serine/threonine-protein kinase 24 isoform a precursor Homo_sapiens_MST3 5
**MDSRAQLWGLALNKRRATLPHPGGSTNLKADPEELFTKLEKIGKGSFGEVFKGIDNRTQKVVAIKIIDLE**
**EAEDEIEDIQQEITVLSQCDSPYVTKYYGSYLKDTKLWIIMEYLGGGSALDLLEPGPLDETQIATILREI**
**LKGLDYLHSEKKIHRDIKAANVLLSEHGEVKLADFGVAGQLTDTQIKRNTFVGTPFWMAPEVIKQSAYDS**
**KADIWSLGITAIELARGEPPHSELHPMKVLFLIPKNNPPTLEGNYSKPLKEFVEACLNKEPSFRPTAKEL**
**LKHKFILRNAKKTSYLTELIDRYKRWKAEQSHDDSSSEDSDAETD**GQASGGSDSGDWIFTIREKDPKNLE
NGALQPSDLDRNKMKDIPKRPFSQCLSTIISPLFAELKEKSQACGGNLGSIEELRGAIYLAEEACPGISD
TMVAQLVQRLQRYSLSGGGTSSH

Green highlight is NLS 278-292 and a possible nuclear export signal NES is in the C-terminal part 335-385 (Lee et al. 2004) behind at least one good, predicted caspase cleavage site. Green highlighted is the part of the NES containg domain that is well conserved in all three sequences.

Probability of caspase cleavage: 0.948435 Site:SDAETD|G (D325)

**Experimentally verified**

```
Probability of caspase cleavage: 0.903749 Site:GSDSGD|W (D336)
```

We reasoned that if we aligned the proteins' truncated N-terminal part that has the NLS signal, the NLS signals in all 3 proteins should align well for the NLS signal  and they do and also for the kinase part that precedes it.

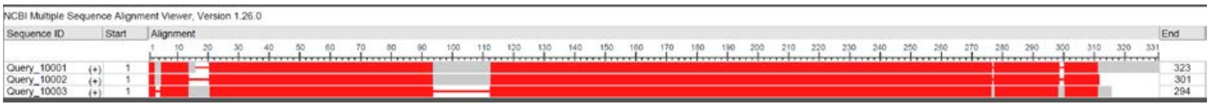

```
Query_10001  1       FGTVRLVGDRGGIVNRPGL-NILDENATNARASRPLESEEDYGEQR-REASPMKRdFALQPLE--TVKapnSRQSS  72
Query_10002  1   [25]FGTVRLVGERGHVVHRPGLlNPLDEAATNARATRSHETEFEQNRQRyRSESPTKR-LSQQQLDndTVK---SRQVS  97
Query_10003          ----------------------------------------------------------------------------

Query_10001  73  PQRKAVPQSQYQPPLSPTRALPQTPLKLPA[6]PRPLRSVHAPvpeSPEYDRELQNELQRDIGMLNLG--YDP   KESR  150
Query_10002  98  PQRRPVPSSPGKTTVQKIEREPETPRAVLK   PAPISFDSAP---SPDYDRELQKQLQREMDGMNIGdmIDP[4]RTGL  172
Query_10003      ------------------------------   ----------------------------------------   ----

Query_10001  151 SPQPQP-SQRPPAPPAHLT[6]QMSLPEIPPYRPSQAVSQQRL--PSLQHASTPQV   LVPLP---GGAQGPRPLPSKN  224
Query_10002  173 IPKTGPaSSQAQKPIAHLG   PIKLPEIPPFRGKHAGSSQPAaqPQTRQYQPPQP[7]LPPHPPYRARDDTPEPPQHQQ  253
Query_10003  1   -------------------   ----------------------------------   ----GQASGGSDSGDWIFTIR  17

Query_10001  225 SPSPSDFTAPSA----------   FPTPSPANPNGELDALNDVIFPALEEALKRRQINLQQVYKP   -GQNPAQIPpQ  287
Query_10002  254 LPTPRPLQTQALQYQQQQQQQP[7]FPSPAPAPPNGEMDALNDVIFPALEEALKRRQIRLQQIQA[9]AGGAPAPTP-R  342
Query_10003  18  EKDPKNLENGALQPSDLDRNKM   --KDIPKRPFSQ--CLSTIISPLFAE--------LKEKSQA   CGGNLGSI---  76

Query_10001  288 QQRAEAAHEKLRKLVYKLAHVCKEIDHYDKAEPVGM--GREVG-SFLEGLLEEILVRVEPLDEDEDvqs  353
Query_10002  343 QQRAEAAHEKIRKLVYRVAHMCKEIDHYDKSEPVHMDTGKDAGGTFLEGFLEEIMLRVEPFDEGEG---  408
Query_10003  77  --------EELRGAIYLAEEACPGISDTMVAQLVQRLQRYSLSGGGTSSH-------------------  118
```

**More detailed alignment showed that the NLSs found in Homo marked and putative in Fg and Mo marked found by alignment of the conserved nuclear active MST3 N-terminal part of MST3 in front of the first predicted or experimentally determined (Homo) caspase cleavage sites.**

```
CLUSTAL multiple sequence alignment by MUSCLE (3.8)


NP_001273578.1      --MAHSPVQSGLPGMQNLKADPEELFTKLEKIGKGSFGEVFKGIDNRTQKVVAIKIIDLE
XP_011324316.1      MADREHDIEEGNEAL-----DPELLYSKEYCIGGGSFGKVYKGVDKRTGQSVAIKVIDIE
XP_003719239.1      MGDRDRDYEEGSP-------DPESLYTKEYCIGGGSFGKVYKGVDKRTGHAVAIKIIDIE
                            :.*         *** *::*   ** ****:*:**:*:** : ****:**:*

NP_001273578.1      EAEDEIEDIQQEITVLSQCDSPYVTKYYGSYLK-------------------LEPGPLDE
XP_011324316.1      SAEDEVEDIIQEIAILSELQSPYVTKYYGSYAKGAELWIVMEFCSGGSCADLMKPGLISE
XP_003719239.1      SAEDEVEDIIQEIAILSELQSPYVTKYYGSYAKGAELWIVMEFCSGGSCADLMKPGLIGE
                    .****:*** ***::**: :*********** *                   ::** :.*

NP_001273578.1      TQIATILREILKGLDYLHSEKKIHRDIKAANVLLSEHGEVKLADFGVAGQLTDTQIKRNT
XP_011324316.1      DYIAIIVRELLMGLDYLHTDKKLHRDVKAANVLLSSNGQVKLADFGVSGQLSATMTKKNT
XP_003719239.1      EYIAIVVRELLLGLDYLHADKKLHRDVKAANVLLAANGQVKLADFGVSGQLSATMTKKNT
                      ** ::**:* ******::**:***:*******: :*:********:***: *  *.**

NP_001273578.1      FVGTPFWMAPEVIKQSAYDSKADIWSLGITAIELARGEPPHSELHPMKVLFLIPKNNPPT
XP_011324316.1      FVGTPFWMAPEVIKQSGYDHKADIWSLGITALELANGEPPYADIHPMKVLFLIPKNPPPR
XP_003719239.1      FVGTPFWMAPEVIKQSGYDHKADIWSLGITALELANGEPPYADIHPMKVLFLIPKNPPPR
                    ****************.** ***********:***.****::::************ **

NP_001273578.1      LEGNYSKPLKEFVEACLNKEPSFRPTAKELLKHKFILRNAKKTSYLTELIDRYKRWKAEQ
XP_011324316.1      LEGNFTKAFKDFIESCLQRDPKDRPTAKDLLRHPFI-RRAKRTTYLTELIERHSRWAAAH
XP_003719239.1      LEGNFTKAFKDFIELCLQRDPKDRPSARELLRHPFV-RHAKKTSYLTELIERHSRWSSTH
                    ****::*.:*:*:* **:.:*. **:*.:**.* *: *.**.*:******:*:.** : :

NP_001273578.1      SHDDSSSEDSDAETD--------------
XP_011324316.1      KGEDDDNWESVNDGRPPAEHERVDEDMWD
XP_003719239.1      KSENDEELD--------------------
                    . ::... :
```

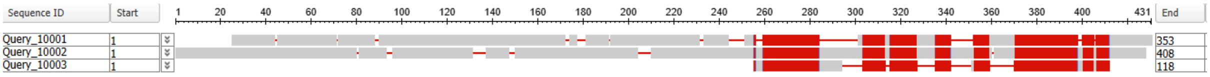


| Accession | Description |
|---|---|
| lcl\|Query_10001 | XP_011324316.1 serine/threonine-protein kinase 24 [Fusarium graminearum PH-1] FGSG_05734 With MST3 domain |
| lcl\|Query_10002 | XP_003719239.1 STE/STE20/YSK protein kinase [Pyricularia oryzae 70-15] MGG_08746 |
| lcl\|Query_10003 | NP_003567.2 serine/threonine-protein kinase 24 isoform a precursor [Homo sapiens] MST3 |
